# Bcl-xL is a key mediator of apoptosis following KRAS^G12C^ inhibition in *KRAS*^*G12C*^ mutant colorectal cancer

**DOI:** 10.1101/2022.05.10.491367

**Authors:** Hajrah Khawaja, Rebecca Briggs, Cheryl Latimer, Md A.M.B. Rassel, Daryl Griffin, Lyndsey Hanson, Alberto Bardelli, Frederica Di Nicolantonio, Simon McDade, Christopher J. Scott, Shauna Lambe, Manisha Maurya, Andreas Lindner, Jochen H.M. Prehn, Jose Sousa, Chris Winnington, Melissa J. LaBonte, Sarah Ross, Sandra Van Schaeybroeck

**Affiliations:** Drug Resistance Group, Patrick G. Johnston Centre for Cancer Research, School of Medicine, Dentistry and Biomedical Science, Queen’s University Belfast, 97 Lisburn Road, Belfast, BT9 7AE, United Kingdom; AstraZeneca, Cambridge, CB2 0RE, United Kingdom; Department of Oncology, University of Torino, Candiolo (TO) 10060, Italy; Candiolo Cancer Institute, FPO-IRCCS, Candiolo (TO) 10060, Italy; Precision Medicine Centre of Excellence, Health Sciences Building, Queen’s University Belfast, 97 Lisburn Road, Belfast, BT9 7AE, United Kingdom; Centre of Systems Medicine; Royal College of Surgeons in Ireland University of Medicine and Health Sciences, 123□St. Stephen’s Green, Dublin 2, Ireland; Personal Health Data Science Group, Sano. Centre for Computational Personalised Medicine, Krakow, Poland

**Author notes:** To whom correspondence should be addressed, at the Patrick G. Johnston Centre for Cancer Research, Queen’s University Belfast, 97 Lisburn Road, Belfast BT9 7AE, Northern Ireland.

**Keywords:** KRAS^G12C^ mutant colorectal cancer, Bcl-xL, KRAS^G12C^ inhibition

## Abstract

**Purpose:** Novel covalent inhibitors of KRAS^G12C^ have shown limited response rates in *KRAS*^G12C^ mutant (MT) colorectal cancer (CRC) patients. Thus, novel KRAS^G12C^ inhibitor combination strategies that can achieve deep and durable responses are needed.

**Experimental design:** Small molecule *KRAS*^G12C^ inhibitors AZ’1569 and AZ’8037 were employed. To identify novel candidate combination strategies for AZ’1569, we performed RNA sequencing, siRNA and high-throughput drug screening. Top hits were validated in a panel of *KRAS*^G12C^MT CRC cells and *in vivo*. AZ’1569-resistant CRC cells were generated and characterised.

**Results:** Response to AZ’1569 was heterogeneous across the *KRAS*^G12C^MT models. AZ’1569 was ineffective at inducing apoptosis when used as single-agent or combined with chemotherapy or agents targeting the EGFR/KRAS/AKT axis. Using a systems biology approach, we identified the anti-apoptotic BH3-family member *BCL2L1*/Bcl-xL as a top hit mediating resistance to AZ’1569. Further analyses identified acute increases in the pro-apoptotic protein BIM following AZ’1569 treatment. ABT-263 (Navitoclax), a pharmacological Bcl-2 family-inhibitor that blocks the ability of Bcl-xL to bind and inhibit BIM, led to dramatic and universal apoptosis when combined with AZ’1569. Furthermore, this combination also resulted in dramatically attenuated tumour growth in *KRAS*^G12C^MT xenografts. Finally, AZ’1569-resistant cells showed amplification of KRAS^G12C^, EphA2/c-MET activation, increased pro-inflammatory chemokine profile and cross-resistance to several targeted agents. Importantly, KRAS amplification and AZ’1569-resistance were reversible upon drug withdrawal, arguing strongly for the use of drug holidays in the case of KRAS amplification.

**Conclusions:** Combinatorial targeting of Bcl-xL and KRAS^G12C^ is highly effective, suggesting a novel therapeutic strategy for *KRAS* ^G12C^MT CRC patients.

## INTRODUCTION

In colorectal cancer (CRC), *KRAS* is the most mutated *RAS* isoform (∼86%), and mutations are most likely to occur in codon 12 (1, 2). KRAS cycles between its inactive, GDP-bound and an active, GTP-bound form that is regulated by either GTP-loading guanine nucleotide-exchange factors or GTPase-activating proteins (GAP) (3). *KRAS* mutations interfere with the rate of its intrinsic and GAP-induced GTP hydrolysis, favouring formation of the constitutively active GTP-bound form.

While substantial advances have been made in the treatment of genetically defined subtypes, such as *RAS/BRAF* wild type (4) and *BRAF*MT CRC (5), an effective therapeutic strategy for *KRAS*MT CRC, the most common genetically defined subtype (∼40-45%) is still lacking. Current treatment options for *KRAS*MT CRC are primarily based on combinations of chemotherapy with antiangiogenic agents (6). KRAS proteins have long been considered “undruggable” due to its small size and the tight binding of KRAS to GTP in its active state. Recently, unique characteristics of the KRAS^G12C^ allele have been exploited for the design of a number of covalent inhibitors that bind specifically to the cysteine at position 12, thereby locking KRAS in its inactive state (7). *KRAS*^G12C^ can be found in ∼14% and ∼4% of lung cancer and CRC respectively (8, 9). A recent trial with the KRAS^G12C^ inhibitor AMG510 (Sotorasib) has shown remarkable single-agent activity in *KRAS*^G12C^MT lung cancer, but efficacy was not encouraging in *KRAS*^*G12C*^MT CRC (10).

Here, we characterize the activity of AZ’1569 (11), a novel KRAS^G12C^ inhibitor, in a panel of *KRAS*^G12C^MT CRC cells. Using RNA sequencing, RNAi/compound screens and mechanistic studies, we identified Bcl-xL as mediator of apoptosis and intrinsic resistance to AZ’1569. We also show that concomitant inhibition of Bcl-xL and KRAS^G12C^ leads to marked increases in therapeutic efficacy in *KRAS*^G12C^MT *in vitro* and xenograft models. Using genomic and proteomic analyses, we show that AZ’1569-acquired resistant models have high level amplification of the KRAS^G12C^ allele with marked elevation of pro-inflammatory environment, resulting in resistance to several targeted agents and conventional chemotherapy.

## MATERIALS AND METHODS

### Materials

AZ’1569 (compound-43), AZ’8037 (compound-25) (11), AZD1480 and AZD6244 (Selumetinib/ARRY-142866) were obtained from Astra Zeneca, 5-Fluorouracil and Oxaliplatin from the Belfast City Hospital Pharmacy (UK), cetuximab from Merck Serono (Middlesex, UK) and crizotinib from Pfizer (Surrey, UK). Ruloxitinib, Capivasertib, ABT-737 and compound library were purchased from Selleckchem (Houston, USA), S6K-18 and ABT-263 from Adooq Biosciences (CA, USA) and SN-38 from Abatra (Shaanxi, China). siRNA targeting *BCL2L1* and the ON-TARGETplus siRNA library was obtained from Dharmacon (Lafayette, USA). See Supplementary methods for details of plasmids.

### Cell culture

C106, SW837, SW1463, SNU1411, LIM2099 and V481 CRC cells were kindly provided by Prof. Bardelli (12). RW7213 cells were provided by Dr. Arango (University hospital Vall d’Hebron, Spain) (13). HCT116 cells were purchased as authenticated stocks from ATCC. Frozen stocks were immediately established from early passage cells. Cells were cultured for not more than 20 passages following thawing. All cell lines were screened monthly for Mycoplasma (MycoAlert Detection Kit, Lonza). KRAS^G12C^ status was confirmed using Sanger sequencing. See Supplementary methods for detailed protocols.

### Generation of AZ’1569-resistant cells

Concentration of AZ’1569 was increased (0.125µM-1µM) until a single-cell density was obtained. Surviving RW7213 cells were expanded in the presence of AZ’1569 (maximum concentration 3µM). Resistance was determined using cell viability assays.

### Protein analysis and Western blotting

Western blotting has previously been described (14, 15). β-actin was used as loading control. Details of antibodies are provided in Supplementary Table S1. Absolute protein quantification of BAK, BAX, BCL2, Bcl-xL and MCL-1 was performed as previously described (16).

### Caspase-3/7 activity assays

Caspase-Glo^®^3/7 activity assays (Promega) have been described previously (15).

### Cell Viability assay

Cell viability was determined using 3-(4,5-dimethylthiazol-2-yl)-2,5-diphenyltetrazolium bromide (MTT) and CellTiter-Glo^®^ (CTG) assays (14), according to the manufacturer’s instructions. IC_50_ values were calculated using the GraphPad Prism 8 (GraphPad Software, Inc.).

### siRNA and DNA transfections

siRNA and DNA transfections were performed using HiPerfect (Qiagen) and X-tremeGENE™ (Merck) respectively, previously described (14).

### RNA/DNA extraction and Real-time reverse transcription-PCR analysis

RNA and DNA extractions were performed using RNeasy^®^ and DNeasy^®^ Blood and Tissue Kits (Qiagen, UK). A260/280 and 260/230 ratios were utilised for quality control. RT-PCR was performed as previously described (15). Probes were purchased from Roche and Thermo Fisher Scientific (TFS, Northants, UK). See Supplementary methods for primer sequences.

### Cytokine/Receptor Tyrosine Kinase (RTK) arrays and CXCL1/TGF-α ELISA

Cytokine, RTK arrays and ELISAs (R&D systems) were used according to the manufacturer’s instructions, as previously described (14). Densitometry was performed using ImageJ.

### RAS-GTP assay

KRAS-GTP expression was evaluated using an active RAS-pull down kit (TFS) according to the manufacturer’s instructions.

### RNA-sequencing

RNA-sequencing of AZ’1569-treated SW837 and SNU1411 cells was performed on a NextSeq 500 using a 150 cycle High Output kit (Illumina, USA) as previously described (14). Additional information is provided in the Supplementary methods.

### Sanger Sequencing

PCR products were cleaned up using Agencourt AM pure beads on the Hamilton Microlab STAR liquid handling robot and Sanger sequencing performed. Electrophoresis of sequencing products was performed on the ABI-3730 48-capillary DNA analyser. Chromatograms were visualised using Geneious software. See Supplementary methods for primer sequences.

### MedExome sequencing and Next Generation Sequencing (NGS)

MedExome and NGS sequencing of RW7213 parental and resistant cells was performed using the Illumina Novaseq 6000 and NextSeq 500 (Illumina, USA) respectively. Additional information is provided in the Supplementary methods.

### *In vitro* migration assays

Migration assays have previously been described (17). Additional information is provided in the Supplementary methods.

### *In vivo* study

*In vivo* studies were conducted as previously described using 6-8-week-old, female NOD SCID mice (Envigo, UK) (14). Details of the initial xenograft growth curves and tolerability studies are provided in the Supplementary methods (Supplementary Fig. S5A-S5B). For the efficacy study, 2.5×10^6^ SNU1411 or 10×10^6^ SW1463 cells were injected into the flank of NOD SCID mice. Mice received vehicle [10% ethanol, 30% polyethylene glycol 400, and 60% Phosal 50 PG orally (PO)], AZ’8037 (100mg/kg PO), Navitoclax (100mg/kg PO) or AZ’8037 (100mg/kg) with Navitoclax (100mg/kg). Each treatment group contained 8 animals. AZ’8037 was administered daily and Navitoclax 5/6 days. Experiments were carried out according to UKCCCR guidelines under licence PPL2875, in accordance with the Animals (Scientific Procedures) Act, 1986, and approved by the Department of Health, Social Services and Public Safety, Northern Ireland.

### Statistical analysis

Robust z-scores (rZ= median/median absolute deviation) were calculated from cell viability assays. All data was plotted (mean and standard deviation, unless specified otherwise), and analysed using GraphPad Prism 8.0. Significance was defined as p<0.05:*; p<0.01:**; p<0.001:***, with p>0.05 not significant (ns). Experiments are representative of 3 independent repeats unless indicated otherwise. The nature of interaction between AZ’1569 and a second drug was determined by calculating CI values according to the Chou-Talalay method (18), using CalcuSyn (Microsoft Windows). CI values <1, >1 and =1 indicate synergy, antagonism, and additive effects respectively. RI values were used when one compound had minimal/no effect on cell viability. RI values>1 indicate synergy (19).

## RESULTS

### *3KRAS*^G12C^MT CRC cells show differential sensitivity to the KRAS^G12C^ inhibitor AZ’1569

To understand the mechanistic basis for the minor clinical responses to KRAS^G12C^ inhibition in CRC (10), we analysed the effect of AZ’1569 in a panel of 7 *KRAS*^G12C^MT CRC cells. Initially, we validated the KRAS mutational status using Sanger sequencing of exon 2, confirming that 4 cell lines had a homozygous *KRAS*^G12C^ mutation and 3 had a heterozygous *KRAS*^G12C^ mutation (Fig. 1A). Five cell lines were found to have a *TP53* mutation, and the V481 cells, were *PIK3CA*MT (Q546P) with loss of PTEN, confirming the results of previous studies (20-23) (https://web.expasy.org/cellosaurus/). Thus, the genetic background of these models captures some of the heterogeneity of *KRAS*MT CRC observed in tumours (24).

**Figure 1.**
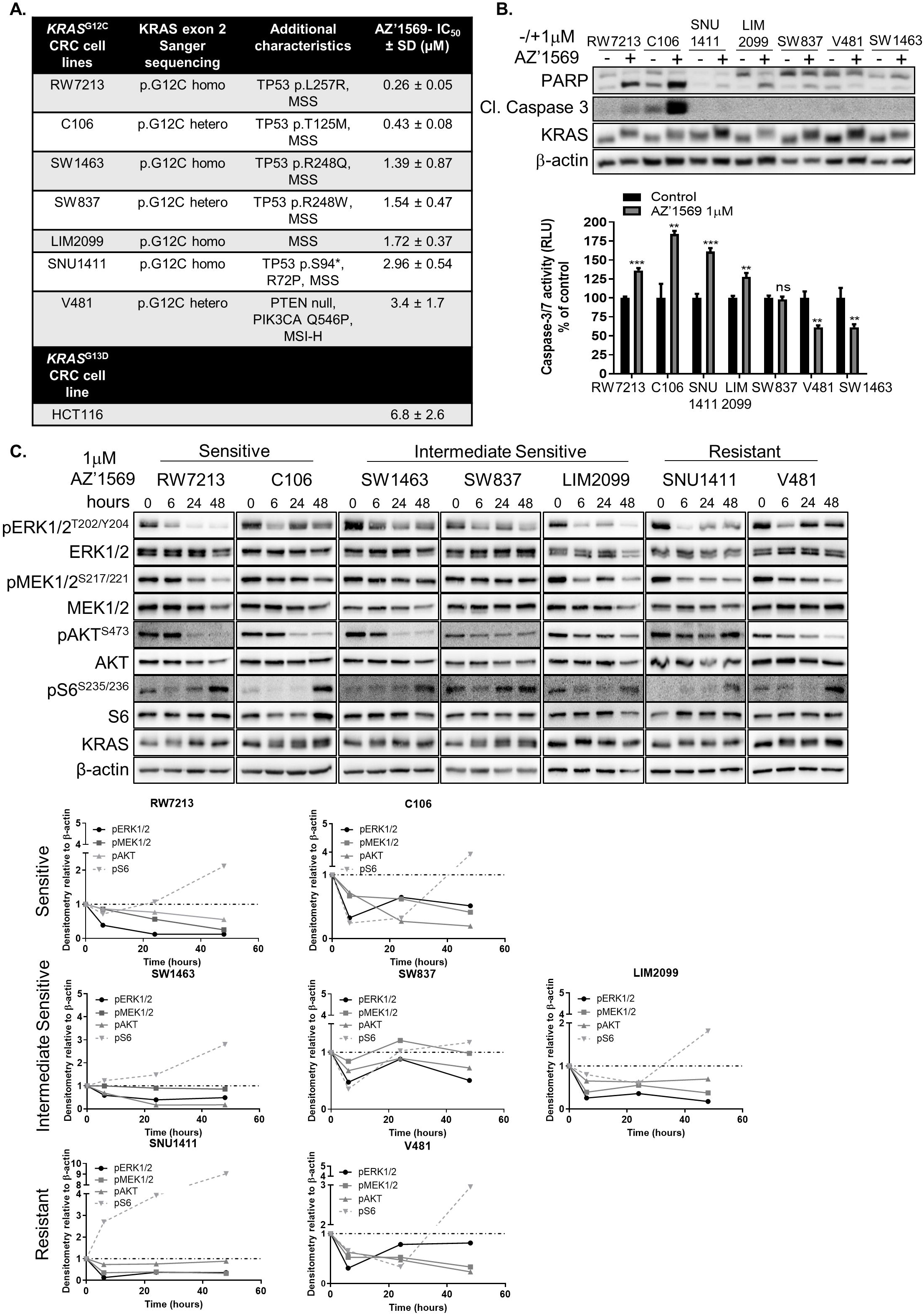
Response to AZ’1569 in *KRAS*^G12C^ MT CRC cells. **A**. KRAS exon 2 Sanger sequencing results for the panel of 7 *KRAS*^G12C^ MT CRC cells. Homozygous (homo), heterozygous (het). Additional mutational changes and MSI (Microsatellite instability) status is also presented (refs. 20-23). CRC cells were treated with AZ’1569 for 120h and cell viability determined using CellTiter-Glo® (CTG) assay. IC_50_ values were calculated using Prism software package. Mean of 3 independent experiments with Standard Deviation is presented in the table. (MSI-H = Microsatellite instability High; MSS = Microsatellite Stable). **B**. CRC cells were treated with AZ’1569 for 48h. PARP, Cleaved C3 and KRAS were determined by Western Blotting (WB) (**Top**), Caspase-3/7 activity levels were measured with values presented as a percentage of their respective controls. Significance was analysed using an unpaired t-test (**bottom**). (Cl = cleaved). **C**. Signaling analysis upon AZ’1569 treatment. *KRAS*^G12C^ MT CRC cell lines were treated with 1µM AZ’1569 for the indicated times and protein lysates were used for WB analysis for the KRAS downstream effectors. Densitometry on WB images was quantified using ImageJ software and normalized to the respective untreated control. Dashed lines on the graphs represent a value of 1.

Response to AZ’1569 was measured using a cell viability assay (Fig. 1A, Supplementary Fig. S1A). RW7213 and C106 cells showed the highest sensitivity to AZ’1569 with IC_50_ values of 0.26µM and 0.43µM, compared to IC_50_ values of 1.39µM, 1.54µM, 1.72µM for SW1463, SW837, LIM2099 cells (“moderately sensitive”) and 2.96 µM and 3.4µM for SNU1411 and V481 cells (“resistant”). Only RW7213 and C106 cells showed marked induction of apoptosis 24h after AZ’1569 treatment, as indicated by PARP cleavage and caspase-3/7 activity (Fig. 1B). Of note, AZ’1569 single-agent activity was not predicted by the G12C zygosity status, p53 mutational status, or by KRAS, EGFR, ERK1/2^T202/Y204^, AKT^S473^ or S6^S235/6^ expression levels (Supplementary Fig. S1A). As expected, *KRAS*^G13D^ MT HCT116 cells were resistant to AZ’1569.

Next, we explored differences in depth, duration and feedback signalling in response to AZ’1569 treatment between our CRC models with low, intermediate and high AZ’1569-IC_50_ values. CRC cells were treated with AZ’1569 in a time course (Fig. 1C). A similar KRAS electromobility shift, indicative of covalent compound binding, was observed following AZ’1569 treatment across the CRC panel, suggesting that differential sensitivity was unlikely due to differences in target engagement (25). KRAS^G12C^ inhibition resulted in profound downregulation in pERK1/2 levels as early as 6h following treatment in all cell lines. Further decreases in pERK1/2 levels were observed in RW7213 and LIM2099 cells 24-48h following treatment, while all the other cell lines showed a rebound in pERK1/2 levels. 24h post-treatment, AZ’1569 caused sustained suppression of pAKT levels in the three most sensitive cell lines, although minor pAKT decreases were also observed in the most resistant cell line, V481. S6 phosphorylation was transiently reduced in the most sensitive cell lines, but marked reactivation was observed in all the cell lines. Altogether, a range of effects of AZ’1569 on downstream signalling dynamics was observed, and these were not sufficient to explain the differences in viability/apoptosis outcome following AZ’1569 treatment.

### KRAS^G12C^ inhibition does not sensitize *KRAS*^G12C^MT CRC cells to chemotherapy

5-FU-based doublet therapies (5-FU+Oxaliplatin; 5-FU+Irinotecan) remains the cornerstone of treatment for *KRAS*MT CRC patients (6). We therefore evaluated whether AZ’1569/5-FU, AZ’1569/Oxaliplatin and AZ’1569/SN-38 combined treatments could effectively suppress the growth of *KRAS*^G12C^MT CRC cells, and used the Chou-Talalay method to calculate combination index (CI) values (18). CI values for combined AZ’1569/5-FU treatment were >0.7 for the majority of the concentrations, indicative of additive interactions (Fig. 2A). Moreover, the absolute cell viability remained > 40% for the majority of combinations. Similar results were obtained for AZ’1569/Oxaliplatin and AZ’1569/SN-38 combinations (Supplementary Fig. S1B). Additionally, except for combined AZ’1569/5-FU treatment in the SNU1411 cells, none of the AZ’1569/chemotherapy combinations resulted in further increases in apoptosis compared to the effect of each treatment alone (Fig. 2B).

**Figure 2.**
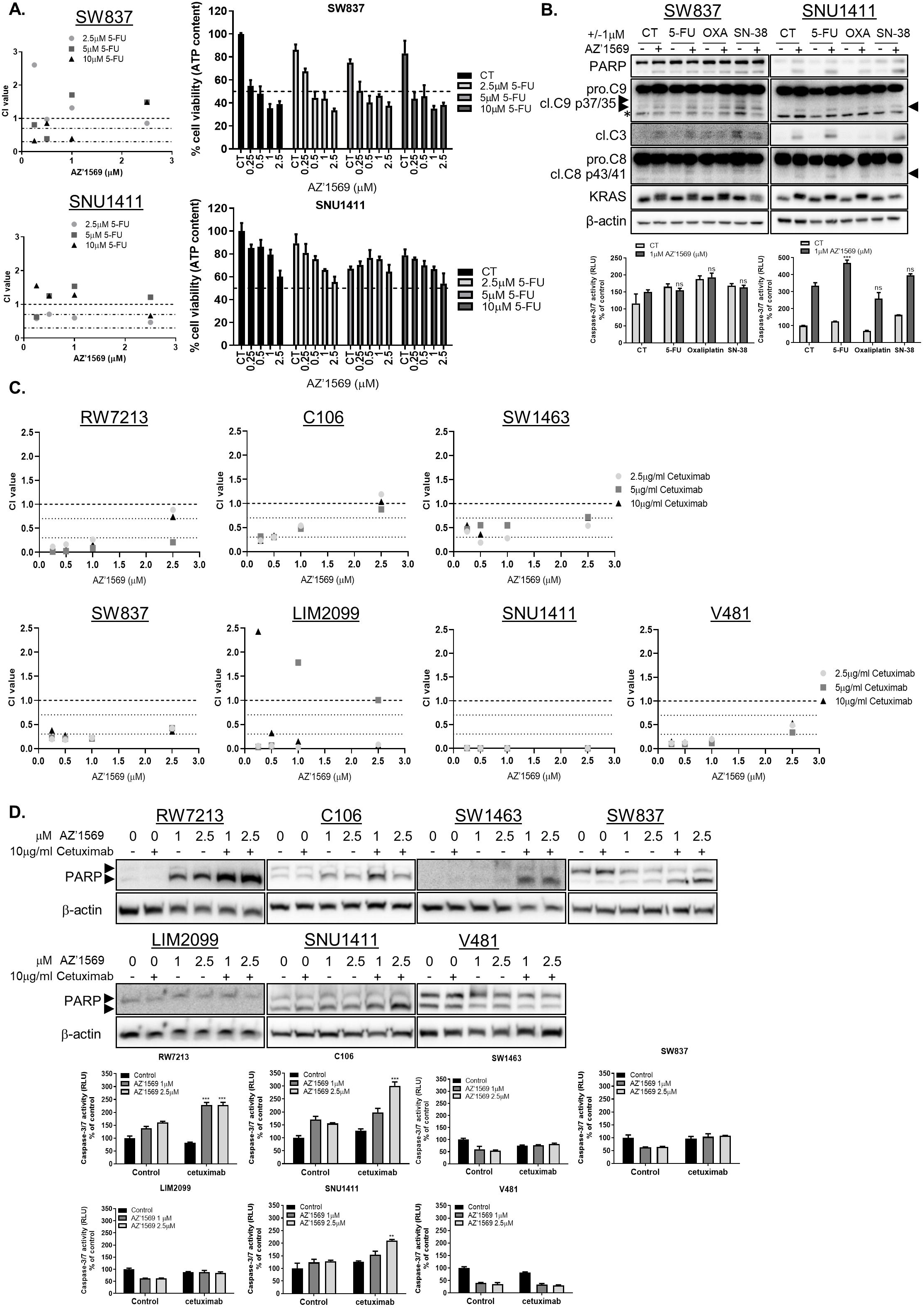
AZ’1569 combined with chemotherapy or Cetuximab does not induce apoptosis in *KRAS*^G12C^ MT CRC cells. **A**. CTG assays in CRC cells treated with no drug (control), 5-FU, AZ’1569 or 5-FU in combination with AZ’1569 for 72h. CI values were calculated using the method of Chou and Talalay. CI values<1, >1, and equal to 1 indicate synergy, antagonism, and additive effects for the drug combinations, respectively. Dashed lines indicate CI values of 0.3, 0.7 and 1. Representative results of at least three experiments (**Left**). Absolute cell viability for different combinations (**right**). Dashed line indicates 50% cell viability. **B**. CRC cells were co-treated with AZ’1569 and 5-FU, Oxaliplatin or SN-38 for 48h. **Upper panel**: PARP, Caspase 9, Caspase 3, caspase 8 and KRAS levels were determined by WB. **Lower panel**: Apoptosis was assessed by Caspase 3/7 activity assay. * = nonspecific band. **C**. CTG cell viability assays in *KRAS*^G12C^ MT CRC cells co-treated with AZ’1569 and Cetuximab for 120h. CI values were calculated to evaluate the nature of interaction. **D**. *KRAS*^G12C^ MT CRC cells were co-treated with AZ’1569 and Cetuximab for 48h, apoptosis was assessed by WB analysis for PARP (**top**) and caspase-3/7 activity assays (**bottom**). A 2-way ANOVA was used to analyse statistical significance.

We next sought to identify pharmacologic combinations that could overcome primary resistance to KRAS^G12C^ inhibition. Based on the known roles for MAPK, AKT, STAT3 and EGFR signalling in intrinsic/acquired resistance to targeted therapies and their relevance in CRC, we next evaluated if combining AZ’1569 with either MEK1/2 inhibitor AZD6244, AKT inhibitor Capivasertib, JAK/STAT inhibitors Ruloxitinib/AZD1480 or EGFR inhibitor cetuximab could effectively suppress CRC cell viability (Figs. 2C-D, Supplementary Figs. S1C-H). Notably, only concurrent cetuximab/AZ’1569 treatment showed moderate/strong synergy across all *KRAS*^G12C^MT CRC cells tested (Fig. 2C). Although combined cetuximab/AZ’1569 treatment resulted in major reduction in cell viability (Supplementary Fig. S1G), efficient apoptosis induction occurred only in RW7213 and C106 cells (Fig. 2D).

### *BCL2L1* is a key regulator of apoptotic response to KRAS^G12C^ inhibition in *KRAS*^G12C^MT CRC cells

Cytostatic and cytotoxic drugs have been linked to clinically observed disease stabilization and objective responses respectively (26). To gain further insight into the molecular mechanisms of apoptosis following AZ’1569 treatment, we performed RNA-seq analysis prior to the onset of cell death in SW837 and SNU1411 cells (Supplementary Figs. S2A-S2B; GSE198530). Significant downregulation of the MAPK pathway-negative feedback mediators *DUSP4/6* and *SPRY4* was observed 6h post-AZ’1569 treatment in both SW837 and SNU1411 cells, confirming inhibition of ERK1/2 activity (Supplementary Fig. S2C). To identify pathways that are involved in resistance to KRAS^G12C^ inhibition, Ingenuity Pathway Analysis (IPA^®^) was conducted using the 3 gene lists generated for both cell lines. IPA^®^ comparison analyses of up- and down regulated pathways showed that 63 pathways overlapped and were significantly deregulated across all the time-points analysed in both cell lines, with a significant enrichment of gene sets in cell death-related signalling pathways, including death receptor, apoptosis, necroptosis and *TP53* signalling (Supplementary Figs. S2D-S2E).

To identify key functional genes/targets that, when inhibited, cooperate with KRAS^G12C^ inhibition to decrease survival, and increase apoptosis in *KRAS*^G12C^MT CRC cells, we used an RNAi screening approach targeting proteins that lie at nodal points in the identified cell death-related signalling pathways. The effect of down-regulating each of these proteins on cell viability was tested in both SW837 and SNU1411 cells, using an ON-TARGETplus siRNA library against 42 targets (Supplementary Fig. S3A) in the absence and presence of AZ’1569 treatment and robust z-score (rZ) values were calculated. Notably, only 1/42 siRNAs had a significant inhibitory effect on survival in the presence of AZ’1569 in both cell lines, and this was *BCL2L1*, the gene encoding the anti-apoptotic BH3-family member Bcl-xL (Fig. 3A). To exclude cell line-specific effects, we extended these studies to a broader panel of *KRAS*^*G12C*^MT CRC cells and also confirmed the cytotoxic activity of the combination by using apoptotic cell death assays, previously described (Fig. 3B). *BCL2L1* silencing resulted in marked increases in apoptosis when combined with AZ’1569 in all *KRAS*^G12C^MT CRC models, compared to the effects of each treatment alone. Additionally, transient overexpression of Myc-tagged Bcl-xL led to marked reduction in basal and AZ’1569-induced apoptosis in SW837 cells (Fig. 3C). Similar effects were observed in the RW7213 cells.

**Figure 3.**
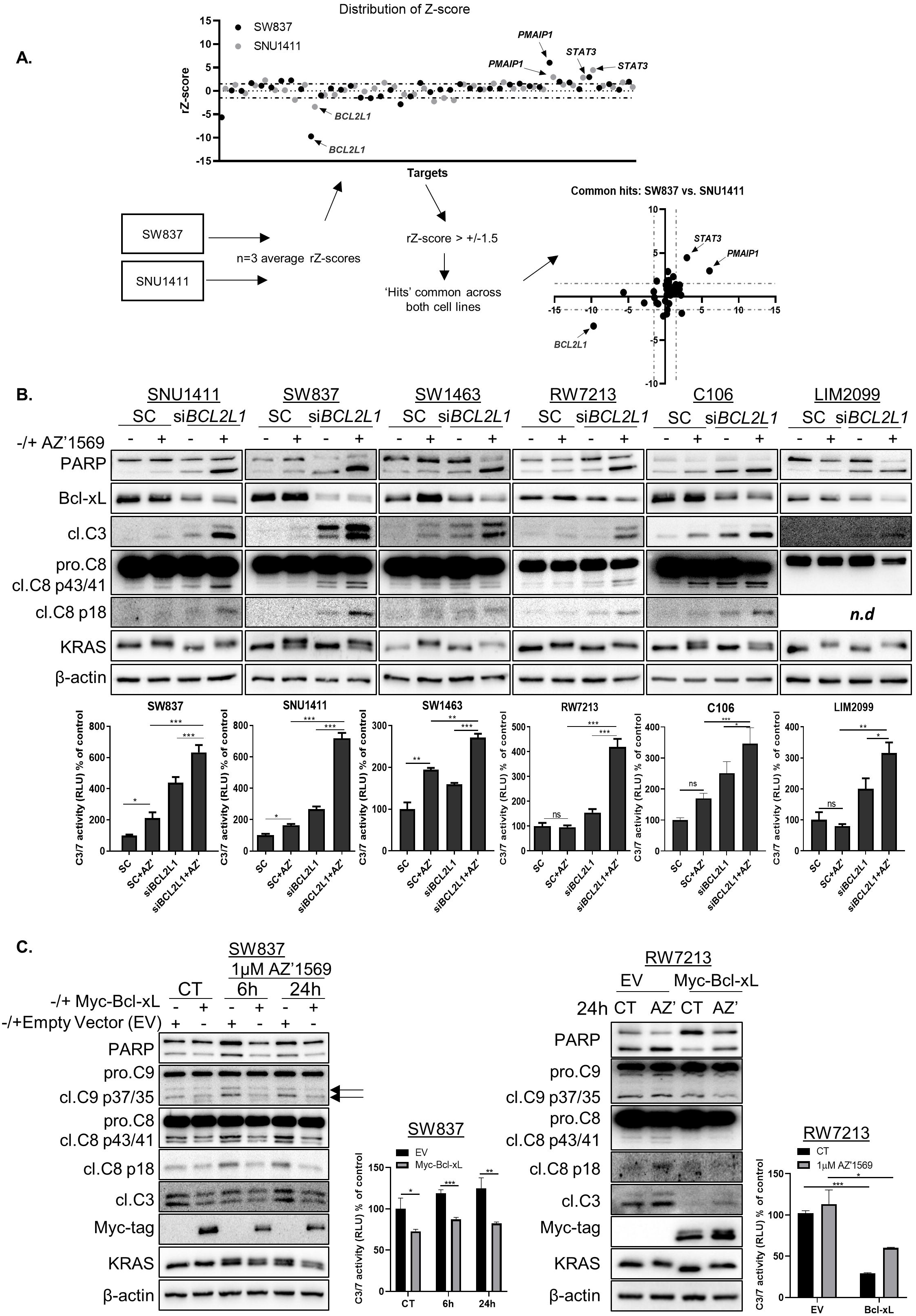
Bcl-xL regulates intrinsic resistance to KRAS^G12C^ inhibition in *KRAS*^G12C^ MT CRC. **A**. Targeted siRNA screen in SW837 and SNU1411 cells. **Top:** SW837 and SNU1411 cells were reverse transfected with 10nM ON-Targetplus siRNA’s targeting 42 genes in the absence or presence of 1µM AZ’1569 for 72h and cell viability was evaluated using the CTG assay. Scatter plot showing robust Z-scores (r-Z) for siRNA screen in SW837 and SNU1411 cells. Positive scores indicate potential mediators of sensitivity to AZ’1569, while negative scores indicate mediators of resistance to AZ’1569. Dashed lines indicate rZ= 0, 1.5 and - 1.5; cut-off thresholds of +/-1.5 were applied to the data. **Bottom**: Schematic of the siRNA approach and analysis. XY graph illustrates hits resulting in sensitisation or resistance to AZD5169 in both cell lines. Data shows average rZ-scores from three independent experiments. **B**. CRC cells were transfected with 10nM On-target SMARTpool siRNA against *BCL2L1* and co-treated with 1µM AZ’1569 (0.25µM AZ’1569 for RW7213 and C106 cells) for 24h (48h for SNU1411) and apoptosis assessed by WB for PARP and Cleaved Caspase 8 and 3 (**Top**) and Caspase 3/7 activity assay (**bottom**). N.d denotes not detected. A 2-way ANOVA was used to evaluate significance. **C**. Expression of PARP, cleaved caspase 9, caspase 8, Myc-tag and KRAS in SW837 and RW7213 cells transiently transfected with 1µg of Myc-tagged Bcl-xL for 24h, followed by treatment with 1µM AZ’1569 (AZD) for the indicated times. Caspase-3/7 activity on cell lysates was also determined. A 2-way ANOVA was used to evaluate significance.

DR_MOMP was previously developed to predict the stress dose required in a cell to induce MOMP (Mitochondrial Outer Membrane Permeabilization), as a read out of sensitivity of CRC cells to genotoxic chemotherapy (16, 27). To evaluate whether sensitivity to AZ’1569 correlated with expression levels of pro- and anti-apoptotic BCL-2 family members or DR_MOMP stress dose, we initially determined the basal absolute BCL-2 proteins profiles (BAK, BAX, BCL-2, Bcl-xL, MCL1) in our *KRAS*^G12C^MT CRC cells (Supplementary Fig. S3B). Not surprisingly, given the heterogeneity of *KRAS*MT CRC, levels of BCL-2 proteins were variable across the cell line panel. Interestingly, Bcl-xL/BAK ratio correlated with response to AZ’1569 (Supplementary Fig. S3B, r=0.54), indicating that cells with an increased Bcl-xL/BAK ratio show an unfavourable response to AZ’1569 treatment. There was no correlation between DR_MOMP calculated stress dose and sensitivity to AZ’1569 treatment. Next, we assessed basal and AZ’1569-induced levels of the pro- and anti-apoptotic BCL-2 family proteins, including the MOMP effector proteins BAX and BAK (Supplementary Fig. S3C). Notably, BIM levels were markedly higher in the AZ’1569-sensitive RW7213 and C106 cells, compared to the levels observed in the intermediate sensitive and resistant cell lines. Expression of BIM was also acutely increased following AZ’1569 treatment in all *KRAS*^G12C^MT CRC cell lines, in particular the RW7213 and C106 cells. Collectively, these data indicate that concomitant suppression of the anti-apoptotic protein Bcl-xL, thereby “freeing” BIM, is needed for a robust apoptotic response following KRAS^G12C^ inhibition in *KRAS*^G12C^MT CRC.

### The BH3-mimetic ABT-737 potently synergizes with KRAS^G12C^ inhibition

To complement the siRNA profiling results, we performed a focused drug screen to identify compounds that could effectively suppress viability of *KRAS*^G12C^MT CRC cells when combined with AZ’1569. We used a drug library targeting the top druggable pathways previously identified (Supplementary Fig. S2D). On the basis of potential for clinical application, we prioritized 45 compounds (Supplementary Fig. S4A), including activators of intrinsic/extrinsic cell death and cell cycle regulators. The effect of these drugs in the absence and presence of AZ’1569 was tested in SW837 and SNU1411 cells. Positive hits were identified as compounds that resulted in robust z-scores less than -1.5 in 3 independent experiments in both cell lines; this identified 12 hits (Fig. 4A). To further refine our hit-list, we determined synergy between these 12 compounds and AZ’1569, using the Chou-Talalay method in SW837 and SNU1411 cells. ABT-737 and Entinostat were the most synergistic with AZ’1569 in both cell lines (Fig. 4B, Supplementary Fig. S4B-C), with ABT-737 resulting in the most potent growth suppression when combined with AZ’1569 in the extended panel of *KRAS*^G12C^MT CRC cells (Fig. 4B, Supplementary Fig. S4D-E). Combined ABT-737/AZ’1569 treatment resulted also in potent increases in apoptosis as indicated by increased PARP cleavage and caspase-9/8/3 processing in all *KRAS*^*G12C*^MT CRC cells (Fig. 4C). Notably, combined ABT-737/AZ’1569 treatment resulted in higher levels of apoptosis, compared to the levels observed with cetuximab/AZ’1569 (Fig. 4D, Supplementary Fig. S4F), suggesting that the ABT-737/AZ’1569 combined strategy could have a more beneficial effect on tumour shrinkage and objective responses in a clinical setting (26).

**Figure 4.**
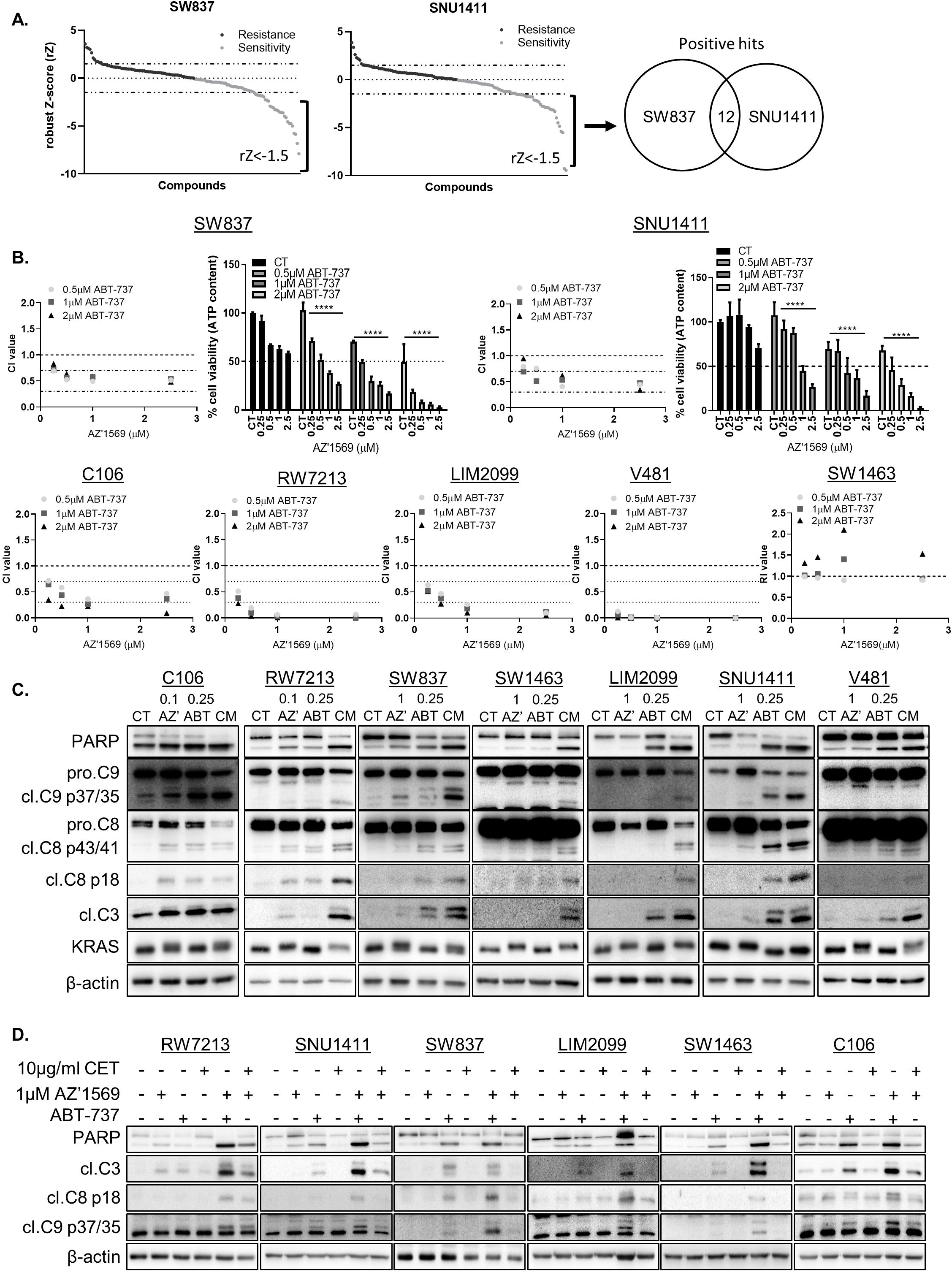
High-throughput drug screen (HTDS) reveals that pharmacological inhibition of Bcl-xL synergizes with KRAS^G12C^ inhibition in *KRAS*^G12C^ MT CRC. **A**. SW837 and SNU1411 cells were co-treated with 1µM AZ’1569 alone or combined with a panel of 45 small molecule inhibitors for 72h and cell viability assessed using the CTG assay. Three concentrations per drug were tested (Supplementary Table S2). Cell viability was analysed using a CTG assay. Scatter plot showing robust Z-scores (r-Z) for each compound concentration used in the drug screen. Negative rZ-scores indicate agents that sensitise to AZ’1569, and *vice versa*. Dashed lines on graphs indicate values of 1.5 and -1.5. Venn diagram indicates number of compounds (past a threshold of r-Z= -1.5), that resulted in sensitization to AZ’1569 in both cell lines. **B**. CTG cell viability assays in *KRAS*^G12C^ MT CRC cells co-treated with AZ’1569 and ABT-737 for 72h. CI values were calculated to evaluate the nature of interaction. Absolute cell viability for AZ’1569/ABT-737 combinations in SW847 and SNU1411 cell lines are also shown. Dashed lines on graphs represent 50% cell viability. **C**. PARP, cleaved C9, cleaved C8, cleaved C3 and KRAS expression levels in *KRAS*^G12C^ MT CRC cells co-treated with AZ’1569 and ABT-737 for 48h (24h for RW7213, SW1463 and V481 cells). CM= combination. **D**. *KRAS*^G12C^ MT CRC cells were treated with AZ’1569 alone or combined with cetuximab or ABT-737 (0.25µM for C106, LIM2099, SNU1411; 0.5µM for SW837, V481, RW7213 and 2.5µM for SW1463) for 48h (24h for C106 cells) and PARP, cleaved C9, cleaved C8 and cleaved C3 determined by WB.

### *KRAS*^G12C^MT CRC xenograft models are sensitive to combinatorial Bcl-xL/KRAS^G12C^ inhibition

We next assessed the *in vivo* therapeutic efficacy of combined Bcl-xL/KRAS^G12C^ inhibition. We selected two different *KRAS*^G12C^MT CRC models, SW1463 and SNU1411 that showed exponential growth characteristics when grown as xenografts (Supplementary Fig. S5A) and used the orally bioavailable BH3-mimetic Navitoclax and the orally bioavailable KRAS^G12C^ inhibitor and close analogue of AZ’1569, AZ’8037. The SW1463 model was resistant to single-agent Navitoclax treatment, and exhibited slowed but persistent growth when mice were treated with AZ’8037 (Fig. 5A). Combination treatment of Navitoclax/AZ’8037 resulted in marked tumour shrinkage in the treated animals. Strong pERK1/2 inhibition was observed, in particular in the Navitoclax/AZ’8037 co-treated tumour samples. Similar to our results in the SW1463 model, single-agent AZ’8037 slowed SNU1411 tumour growth (Fig. 5B). There was also no effect of single-agent Navitoclax. Although addition of Navitoclax to AZ’8037 resulted in further reduction in tumour growth, there was no tumour regression in the SNU1411 xenografts. Treatment cessation resulted in tumour regrowth in AZ’8037 monotherapy and Navitoclax/AZ’8037 combination groups (Fig. 5B). The Navitoclax/AZ’8037 combination was less well tolerated in this second mouse model as shown by decreases in tumour weight in week 2 of the treatment (Fig. 5B). Navitoclax was therefore given as a 5 day on, 1 day off schedule. Collectively, these results indicate that Bcl-xL-targeted agents may be highly effective when used in combination with KRAS^G12C^ inhibitors in *KRAS*^G12C^MT CRC.

**Figure 5.**
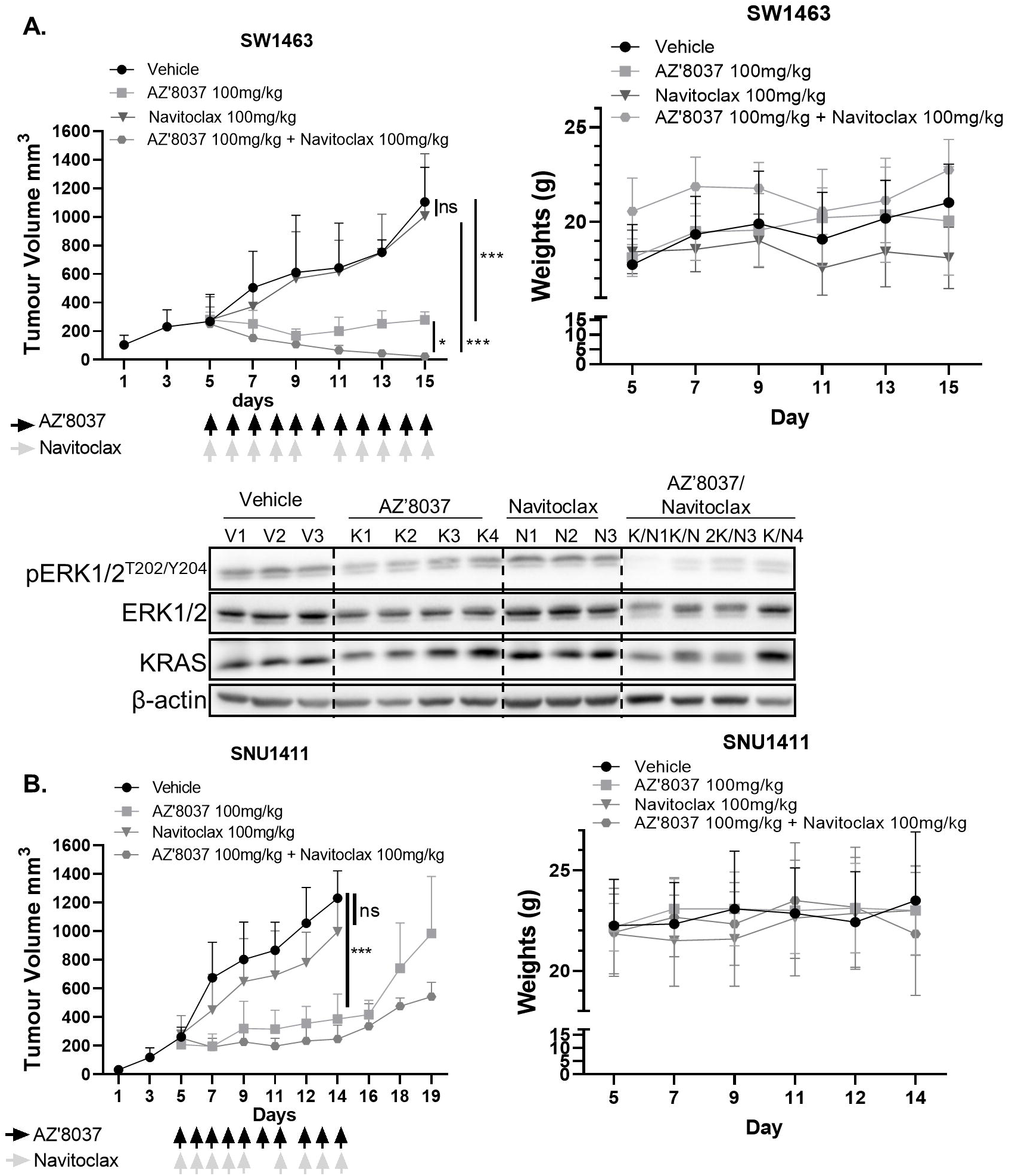
Combined KRAS^G12C^ and Bcl-xL inhibition results in reduction in growth of *KRAS*^G12C^ MT CRC *in vivo*. **A**. Growth rate (**left**) and mouse weight (**right**) of SW1463 xenografts in NOD/SCID mice treated with vehicle, AZ’8037, Navitoclax, or AZ’8037 in combination with Navitoclax. Differences in growth were determined using a one-way ANOVA with Tukey’s test for multiple comparisons. WB analysis for pERK1/2, ERK1/2 and KRAS in tumour samples collected at day 15. **B**. Growth rate (**left**) and mouse weight (**right**) of SNU1411 xenografts in NOD/SCID mice treated with vehicle, AZ’8037, Navitoclax, or AZ’8037 in combination with Navitoclax.

### Generation of CRC models with acquired resistance to KRAS^G12C^ inhibition

Although recent clinical trials of KRAS^G12C^ inhibition have shown modest efficacy in *KRAS*^G12C^MT CRC, emergence of acquired resistance limits further clinical benefit (28). In order to identify mechanisms underlying acquired drug-resistance to KRAS^G12C^ inhibition and therapeutic strategies to overcome this limitation, we generated a preclinical AZ’1569-resistant CRC model. We selected the RW7213 cell line, which shows the highest sensitivity to AZ’1569, and cultured this cell line until resistant derivatives/clones emerged in the presence of AZ’1569. Three independent resistant (R) RW7213 cell populations were obtained and these were therefore indicated as resistant #2, #3 and #4 (Supplementary Fig. S6A). Resistance to AZ’1569 was confirmed by cell viability assays comparing parental and resistant cell derivatives (Fig. 6A). All resistant models also showed cross-resistance to the KRAS^G12C^ inhibitors Sotorasib and Adagrasib (MRTX849) (Fig. 6A, Supplementary Fig. S6A). Notably, both AZ’1569 and Sotorasib seemed to increase the growth rate of AZ’1569-R clones, suggesting that AZ’1569-R clones had become addicted to the presence of KRAS^G12C^ inhibition for proliferation.

**Figure 6.**
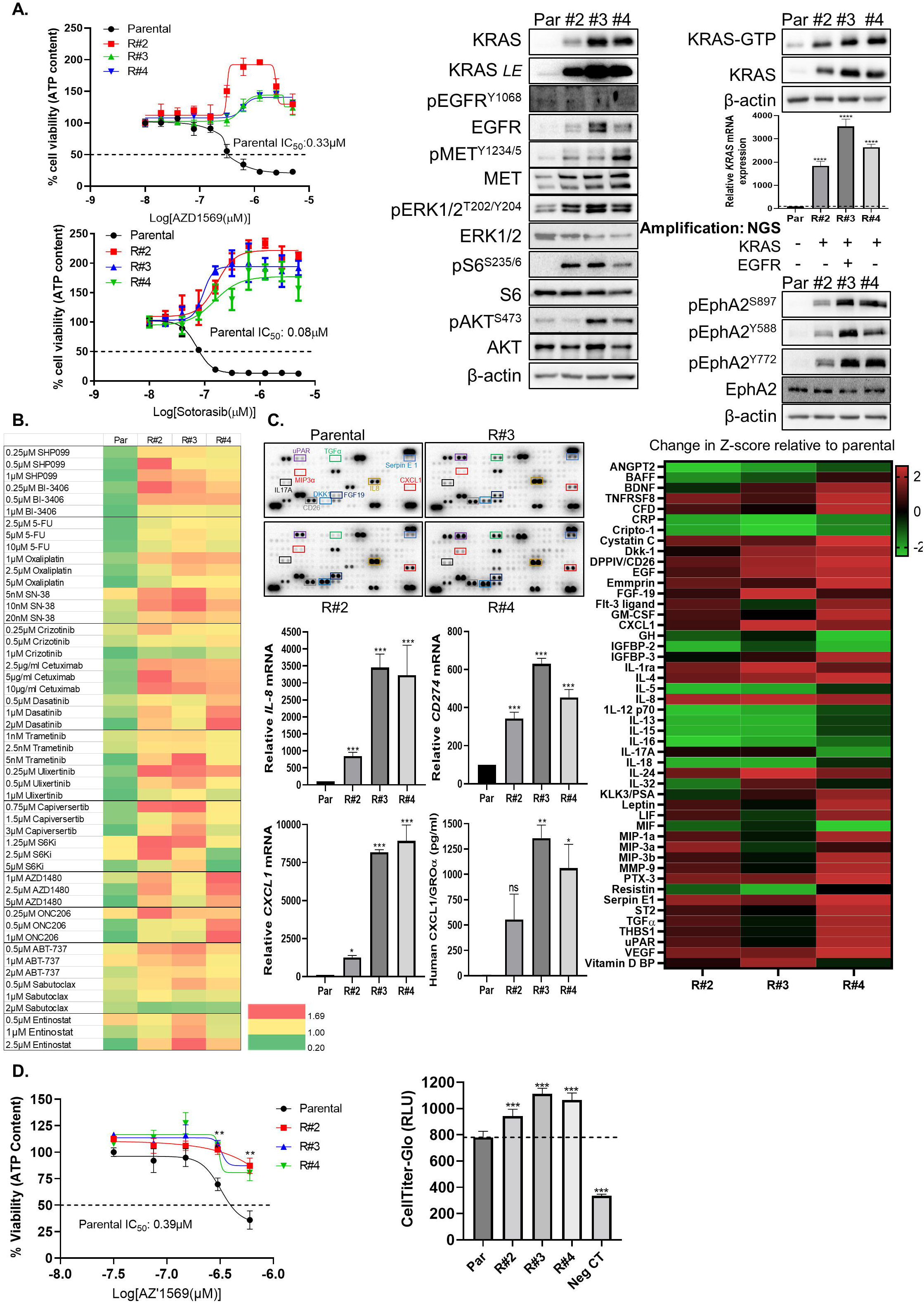
AZ’1569-acquired resistant cells exhibit increased PD-L1 expression and a pro-inflammatory phenotype. **A. Left:** RW7213 parental and AZ’1569 resistant clones (#2, #3 and #4) were treated for 72h with indicated concentrations of AZ’1569 or Sotorasib, and cell viability was determined using CTG assays. **Right:** Lysates from RW7213 parental and AZ’1569 resistant clones were analysed by WB for KRAS, pEGFR^Y1068^, EGFR, pMET^Y1234/1235^, MET, pERK1/2^T202/Y204^, ERK1/2, pS6^S235/6^, S6, pAKT^S473^, AKT, pEphA2^S897^, pEphA2^Y588^, pEphA2^Y772^ and EphA2. Active Raf1-bound Ras was isolated from RW7213 parental and resistant clones using a RAS-GTP assay and basal GTP-bound and total KRAS levels assessed by WB. *LE*=longer exposure. *KRAS* mRNA was quantified using RT-PCR. Raw values were normalised to *ACTB* and *GAPDH* expression and were analysed using the ΔΔCT method. A one-way ANOVA was used to calculate statistical significance. Data is representative of three independent experimental repeats. Results of NGS sequencing of RW7213 Par and AZ’1569-R clones are shown. **B**. RW7213 parental and resistant cells were treated with SHP-099, BI-3406, 5-FU, SN-38, Oxaliplatin, Crizotinib, cetuximab, Dasatinib, Trametinib Ulixertinib, Capivasertib, PF-4708671, AZD1480, ABT-737, Sabutoclax or Entinostat for 72h, at the indicated concentrations and cell viability was assessed using CTG assays. Heatmap represents cell viability relative to control. Data is representative of three independent experimental repeats. **C. Top Left**: Human cytokine array using conditioned medium of RW7213 parental and resistant clones. **Right**: Mean spot pixel density was analysed using Image J, robust Z-scores (r-Z) (relative to parental cells) were calculated using densitometry data and presented in a heatmap. **Bottom Left**: *CXCL1, CD274* and *IL8* mRNA in parental and resistant clones was quantified using RT-PCR. Raw values were normalised to the expression of housekeeping genes *ACTB* and *GAPDH* and were analysed using the ΔΔCT method. CXCL1 protein levels in the culture media of parental (Par) and resistant subpopulations were measured by ELISA. A one-way ANOVA was used to calculate statistical significance. Data is representative of three independent experimental repeats. **D. Left:** Dose response curves for AZ’1569 in RW7213 cells, incubated with conditioned media from parental cells or drug resistant clones #2, #3 or #4. Cells were treated for 72h and cell viability was determined using CTG assay. IC_50_ values were calculated using Prism software package. Dashed line indicates 50% cell viability. A representative of three independent experiments is shown. **Right:** A 24-well 5µm polycarbonate Transwell® insert-plate system was used. 2.5×10^5^ peripheral blood mononuclear cells (PBMCs) were re-suspended in 2% FCS-supplemented DMEM and were added to the top chamber. The bottom chamber was filled with conditioned medium (medium=2% FCS-supplemented DMEM) obtained from RW7213 parental and resistant cells. Cells were incubated for four hours, following which CellTiter-Glo® was used to measure PBMC migration to the bottom chamber (RLU= relative luminescence). Serum-free DMEM was used in the bottom chamber as a negative control (Neg CT). Data is representative of three independent experimental repeats.

### Acquired AZ’1569-R clones display KRAS amplification and activation of several RTKs

Prior studies indicated that tumours with acquired resistance to KRAS^G12C^ agents can have multiple resistance mechanisms, including alterations within the RAS/MAPK pathway and bypass activating alterations (28, 29). Although all resistant cell populations retained the original KRAS^G12C^ mutation, MedExome sequencing of the RW7213-R clones did not reveal any secondary mutations within KRAS, BRAF/MEK/ERK, PIK3CA/AKT (Supplementary Fig. S6B). Notably, total KRAS, KRAS-GTP and mRNA levels together with pERK1/2 levels were markedly upregulated in the RW7213-R clones (Fig. 6A). Moreover, next-generation sequencing confirmed that *KRAS* was amplified in all three clones and clone 3 had an additional amplification in EGFR (Supplementary Fig. S6C). Of note, withdrawal of AZ’1569 for 8 weeks resulted in loss of KRAS over-expression, reversal of hyperactivated signalling to ERK1/2 and re-sensitisation to AZ’1569 (Supplementary Fig. S6D).

We also assessed the phosphorylation status of 49 RTK’s in parental and AZ’1569-R clones. AZ’1569-R derivatives showed increased phosphorylation of a number of RTKs such as c-MET and EphA2 (Supplementary Fig. S6E); these were validated using WB analysis (Fig. 6A). Collectively, our data revealed KRAS amplification and coincidental bypass RTK acquired alterations in our AZ’1569-resistant clones, suggesting that the cell models generated in this work have the potential to recapitulate clinically-relevant resistance mechanisms.

### Acquired AZ’1569-R clones driven by KRAS amplification represents a therapeutic challenge

Classically, the identification of molecular mechanisms underlying acquired resistance should enable rational intervention with small-molecule inhibitors to overcome resistance. Based on the results from our RTK array validation, we used a focused drug screen targeting KRAS/ERK signalling, including SOS1-KRAS inhibitor BI-3406, SHP2 inhibitor SHP-099, MEK1/2 inhibitor Trametinib and ERK1/2 inhibitor Ulixertinib. We also used inhibitors of AKT (Capivasertib), S6K (PF-4708671), JAK/STAT3 (AZD1480), EphA2 (Dasatinib), c-MET (Crizotinib) kinases and BCL-2/Bcl-xL inhibitors ABT-737 and Sabutoclax (Fig. 6B). Bcl-xL inhibition did not affect survival of AZ’1569-R clones (Fig. 6B; Supplementary Fig. S6F). Surprisingly, we found that AZ’1569-R derivatives displayed reduced sensitivity to the diverse kinase inhibitors compared to their parental counterpart. Moreover, AZ’1569-R clones were also more resistant to the chemotherapeutic agents 5-FU, SN-38 and Oxaliplatin. As expected, all AZ’1569-R clones were highly resistant to cetuximab.

### Acquired AZ’1569-R clones overproduce a wide array of pro-inflammatory factors

We have previously shown that oncogenic KRAS regulates growth factor/cytokine shedding and ADAM17 activity (30), a protease involved in acute resistance to chemotherapy and targeted therapies (31). We therefore investigated the growth factors/cytokines released by the AZ’1569-resistant cells, using a cytokine array (Fig. 6C). Of the 105 cytokines examined by the array, 15 targets were >1.5-fold upregulated in all 3 resistant clones, and these included cytokines/chemokines involved in innate/adaptive immunity (eg. IL-8, CXCL1) and growth factors (eg. TGFα). We validated our array results using real-time PCR and/or specific ELISAs, showing that AZ’1569-resistant clones exhibited higher levels of IL-8, CXCL1, IFNγ and TGFα (Fig. 6C, Supplementary Fig. S6G-H). Given the marked cytokine/chemokine abundance in the drug resistant lines, we also determined PD-L1 levels and found >300-fold increased levels of *CD274* (encoding PD-L1) in the AZ’1569-R clones (Fig. 6C). Of note, conditioned medium of all three AZ’1569-R clones markedly reduced sensitivity of parental RW7213 cells to AZ’1569 (Fig. 6D). Additionally, exposure to conditioned medium of all three AZ’1569-R clones increased PBMC migration, indicating their importance for lymphocytic infiltration (Fig. 6D). Taken together, these results show that long-term exposure to AZ’1569 dramatically increases the immunogenicity of these cells and suggests that the induction of pro-inflammatory factors may produce a tumour microenvironment that is conducive to increased tumour infiltration by immune cells.

## DISCUSSION

The efficacy of anti-cancer targeted therapies has often been compromised by the occurrence of intrinsic and acquired resistance mechanisms, involving intra-tumoural heterogeneity and various compensatory signalling. Recently, drugs such as Sotorasib and Adagrasib, which inhibit KRAS^G12C^, have emerged as promising targeted therapies for *KRAS*^G12C^MT patients (32, 33). However, clinical trials, such as CodeBreaK100 and KRYSTAL-1 using single-agent Sotorasib and Adagrasib respectively, have shown substantial differences in response rates between lung cancer and CRC patients (10, 34, 35). On the basis of these studies, Sotorasib was granted FDA-approval, but only for *KRAS*^G12C^MT lung cancer patients (36). Understanding the factors underlying intrinsic/acquired resistance to this new class of compounds is critical, in particular for CRC patient benefit. There is a growing body of evidence suggesting that deficient apoptosis induction following targeted therapy treatments can lead to a lack of efficacy (37, 38). Indeed, we found that KRAS^G12C^ inhibitor monotherapy was relatively ineffective at inducing apoptosis *in vitro* in *KRAS*^G12C^MT CRC.

There are, as of yet, no available predictive biomarkers for response to KRAS^G12C^ inhibitors. In agreement with previous studies, we found that sensitivity to AZ’1569 was not predicted by the KRAS allele zygosity status, the presence of concomitant mutations (including *TP53* mutations) or baseline levels of kinases within the EGFR/KRAS axis (32, 39). We previously employed DR_MOMP, an apoptosis predictor model that requires protein profiling of Bcl-2 family proteins to predict therapeutic response and prognosis in CRC (16, 27). Although not significant, we found that the Bcl-xL/BAK ratio correlated with response to AZ’1569 treatment. Previous *in vitro* and clinical studies have shown positive correlations between BIM expression and response to anti-EGFR and BRAF drugs in EGFR and BRAF addicted tumours (40, 41). Interestingly, our study showed that pre-treatment BIM levels were associated with sensitivity and apoptotic response to AZ’1569 in *KRAS*^G12C^MT CRC cells. Further biomarker analysis of tissue samples from patients treated with KRAS^G12C^ inhibitors will be needed to confirm the predictive role of BIM.

Previous studies evidenced that RTK/kinase feedback activation is among the main mechanisms of adaptive resistance to KRAS^G12C^ inhibitors and that therefore vertical combinations with RTK/SHP2 inhibitors (42) or kinase inhibitors (eg. MEK1/2, PI3K/mTORC1/2) (32, 39, 43) are the most attractive combination options. In support of these studies, we found consistent evidence of rapid ERK1/2, AKT and/or S6 feedback reactivation following AZ’1569 treatment, although this was heterogeneous across the *KRAS*^G12C^MT models. Combinations of AZ’1569 with inhibitors of these feedback loops showed differential effects across the *KRAS*^G12C^MT CRC cell lines. Conversely, inhibition of EGFR markedly enhanced sensitivity to AZ’1569 in all *KRAS*^G12C^MT CRC cells, supporting the findings of a recent study (12). However, addition of cetuximab to AZ’1569 only resulted in potent increases in cell death in 3/7 *KRAS*^G12C^MT CRC models.

Using RNA-sequencing, IPA^®^ pathway analysis and a siRNA screening approach, we identified that *BCL2L1* was a critical mediator of resistance to cell death following KRAS^G12C^ inhibition in CRC cells. Moreover, using focused drug screens, we identified that the BCL-2/Bcl-xL inhibitor ABT-737 was an effective inducer of apoptosis when combined with AZ’1569 in the panel of *KRAS*^G12C^MT CRC cells. Treatment with AZ’1569 resulted in acute increases in the pro-apoptotic protein BIM, which may ‘prime’ cells for death, but was insufficient to cause apoptosis in 5/7 *KRAS*^G12C^MT CRC models due to the presence of inhibitory anti-apoptotic proteins, such as Bcl-xL. Consistent with previous studies, we demonstrated that ABT-737 abrogates the inhibitory complex between Bcl-xL and BIM (Supplementary Fig. S3D) (44), leading to robust increases in apoptosis when ABT-737 wascombined with KRAS^G12C^ inhibition in our study. Alongside its pivotal role in regulating MOMP, Bcl-xL has been identified as a critical mediator of stem cell survival through the adeno-to-colon carcinoma sequence (45). Additionally, a number of studies have shown that Bcl-xL plays an important role in regulating sensitivity to chemotherapy and other targeted therapies (45-47).

The importance of Bcl-xL as a mediator of acute resistance to KRAS^G12C^ inhibitors was demonstrated *in vivo*, where combined treatment of *KRAS*^G12C^MT CRC xenografts with the BCL-2/Bcl-xL and KRAS^G12C^ inhibitors, Navitoclax and AZ’8037, resulted in supra-additive reductions in tumour growth or regression. During this manuscript’s preparation, initial results of the phase Ib study of cetuximab with Adagrasib were released, showing a response rate of 43% in *KRAS*^G12C^MT CRC patients (48). Although initial results of this small study are encouraging, it also suggests that a major part of this population will not respond to this combination, indicating the need for alternative treatment combinations. Our data suggests that combined Bcl-xL/KRAS^G12C^ inhibition is another potential novel treatment strategy for this molecular subgroup of CRC patients. Although less well tolerated in our *in vivo* strain, combination treatments with Navitoclax have been widely trialled in other *in vivo* strains and patients without major reported toxicities (46, 49). In further support of our data, a recent study showed that the Bcl-xL-targeted PROTAC, DT2216, enhanced the therapeutic efficacy of Sotorasib in the SW837 *KRAS*^G12C^MT model, and demonstrated also a good tolerability (50).

Acquired resistance is a major problem limiting clinical efficacy of targeted therapies. We observed amplification of the KRAS^G12C^ allele in all 3 AZ’1569-acquired resistant clones, which also coincided with acquired bypass activations in a number of RTK’s. Interestingly, this is consistent with analysis of clinical samples from patients treated in phase I/II studies with Adagrasib (28). Contrastingly, no acquired mutations affecting the switch II pocket of KRAS (R68S, H95D/Q/R, Y96C) or other pathogenic mutations in other RTK-RAS-MAPK pathway members were detected. Importantly, we also show that acquired resistance driven by KRAS amplification is reversible upon drug withdrawal, likely because KRAS amplification confers a selective disadvantage in the absence of KRAS^G12C^ inhibition.

The AZ’1569-R cells demonstrated a high level of resistance to a range of targeted therapies, particularly SOS1 and MEK/ERK inhibition. Importantly, AZ’1569-R cells also showed markedly reduced sensitivity to the 3 chemotherapies used in CRC treatment. Thus, our results would indicate that, at least in cases where KRAS^G12C^ inhibitor resistance is driven by KRAS amplification, patients who progress following upfront treatment with KRAS^G12C^ inhibition may be poor candidates for other targeted therapies or chemotherapies. A previous study has shown that Sotorasib has an early impact on tumour immune cell infiltration (33). Interestingly, our acquired AZ’1569-R cells showed a markedly increased pro-inflammatory cytokine/chemokine profile, which resulted in increased lymphocytic infiltration. These data further suggest major changes in the immune microenvironment of KRAS^G12C^ inhibitor-resistant tumours, which may affect their response to immune-targeted therapies.

In conclusion, using a systems biology approach, we identified Bcl-xL as an important mediator of intrinsic resistance to KRAS^G12C^ inhibition in *KRAS*^*G12C*^MT CRC. We show that KRAS^G12C^ inhibition primes cells for death through acute induction of BIM, with co-neutralization of Bcl-xL resulting in potent increases in cell death. From a cancer therapeutics perspective, the substantial tumour growth inhibition observed in our xenografts provides a strong rationale to combine Bcl-xL blockade, using Navitoclax or HDAC1-3 inhibition (Supplementary Fig. S4C) with KRAS^G12C^ inhibitors in CRC patients. We also demonstrate the importance of drug holidays, in order to delay/overcome emergent resistance to KRAS^G12C^ inhibition. Finally, cross-resistance to other targeted therapies and importantly conventional chemotherapy in the AZ’1569-R cells poses a challenge, with implications for the optimal use of KRAS^G12C^ inhibitors as a second or third line option.

## Notes

**Financial support:** Cancer Research UK (C212/A7402); MErCuRIC, funded by the European Commission’s Framework Programme 7, under contract #602901; Tom Simms Memorial Fund in Queen’s University Belfast; sponsored research agreement from Astra Zeneca; Science Foundation Ireland and the Health Research Board (16/US/3301).

**Conflicts of interest:** Sarah Ross & Lyndsey Hanson were/are employees and shareholders of AstraZeneca. The other authors declare that they have no conflict of interest.

https://www.ncbi.nlm.nih.gov/geo/query/acc.cgi?acc=GSE198530

https://eur02.safelinks.protection.outlook.com/?url=%3A%2F%2Fwww.ncbi.nlm.nih.gov%2Fbioproject%2F815942&amp;data=04%7C01%7CH.Khawaja%40qub.ac.uk%7Cff0dc5d74e48464dc0ca08da05a17a12%7Ceaab77eab4a549e3a1e8d6dd23a1f286%7C0%7C0%7C637828488513915676%7CUnknown%7CTWFpbGZsb3d8eyJWIjoiMC4wLjAwMDAiLCJQIjoiV2luMzIiLCJBTiI6Ik1haWwiLCJXVCI6Mn0%3D%7C2000&amp;sdata=LvT3AeKHbh7FCsesGA9JE9Jl4nDwrOT8dWLKV1SCkjA%3D&amp;reserved=0

### Competing Interest Statement

Sarah Ross & Lyndsey Hanson were/are employees and shareholders of AstraZeneca. The other authors declare that they have no conflict of interest.

## REFERENCES

1. Cox AD, Fesik SW, Kimmelman AC, Luo J,Der CJ. Drugging the undruggable RAS: Mission possible? Nat Rev Drug Discov 2014; 13: 828–51.

2. Prior IA, Hood FE,Hartley JL. The Frequency of Ras Mutations in Cancer. Cancer Res 2020; 80: 2969–74.

3. Simanshu DK, Nissley DV,McCormick F. RAS Proteins and Their Regulators in Human Disease. Cell 2017; 170: 17–33.

4. Tejpar S, Stintzing S, Ciardiello F, Tabernero J, Van Cutsem E, Beier F, et al. Prognostic and Predictive Relevance of Primary Tumor Location in Patients With RAS Wild-Type Metastatic Colorectal Cancer: Retrospective Analyses of the CRYSTAL and FIRE-3 Trials. JAMA Oncol 2017; 3: 194–201.

5. Kopetz S, Grothey A, Yaeger R, Van Cutsem E, Desai J, Yoshino T, et al. Encorafenib, Binimetinib, and Cetuximab in BRAF V600E-Mutated Colorectal Cancer. N Engl J Med 2019; 381: 1632–43.

6. Kubicka S, Greil R, Andre T, Bennouna J, Sastre J, Van Cutsem E, et al. Bevacizumab plus chemotherapy continued beyond first progression in patients with metastatic colorectal cancer previously treated with bevacizumab plus chemotherapy: ML18147 study KRAS subgroup findings. Ann Oncol 2013; 24: 2342–9.

7. Ostrem JM, Peters U, Sos ML, Wells JA,Shokat KM. K-Ras(G12C) inhibitors allosterically control GTP affinity and effector interactions. Nature 2013; 503: 548–51.

8. Blons H, Emile JF, Le Malicot K, Julie C, Zaanan A, Tabernero J, et al. Prognostic value of KRAS mutations in stage III colon cancer: post hoc analysis of the PETACC8 phase III trial dataset. Ann Oncol 2014; 25: 2378–85.

9. Dunnett-Kane V, Burkitt-Wright E, Blackhall FH, Malliri A, Evans DG,Lindsay CR. Germline and sporadic cancers driven by the RAS pathway: parallels and contrasts. Ann Oncol 2020; 31: 873–83.

10. Hong DS, Fakih MG, Strickler JH, Desai J, Durm GA, Shapiro GI, et al. KRAS(G12C) Inhibition with Sotorasib in Advanced Solid Tumors. N Engl J Med 2020; 383: 1207–17.

11. Kettle JG, Bagal SK, Bickerton S, Bodnarchuk MS, Breed J, Carbajo RJ, et al. Structure-Based Design and Pharmacokinetic Optimization of Covalent Allosteric Inhibitors of the Mutant GTPase KRAS(G12C). Journal of medicinal chemistry 2020; 63: 4468–83.

12. Amodio V, Yaeger R, Arcella P, Cancelliere C, Lamba S, Lorenzato A, et al. EGFR Blockade Reverts Resistance to KRAS(G12C) Inhibition in Colorectal Cancer. Cancer discovery 2020; 10: 1129–39.

13. Tibbetts LM, Chu MY, Vezeridis MP, Miller PG, Tibbetts LL, Poisson MH, et al. Cell culture of the mucinous variant of human colorectal carcinoma. Cancer Res 1988; 48: 3751–9.

14. Van Schaeybroeck S, Kalimutho M, Dunne PD, Carson R, Allen W, Jithesh PV, et al. ADAM17-dependent c-MET-STAT3 signaling mediates resistance to MEK inhibitors in KRAS mutant colorectal cancer. Cell reports 2014; 7: 1940–55.

15. Khawaja H, Campbell A, Roberts JZ, Javadi A, O’Reilly P, McArt D, et al. RALB GTPase: a critical regulator of DR5 expression and TRAIL sensitivity in KRAS mutant colorectal cancer. Cell death & disease 2020; 11: 930.

16. Lindner AU, Concannon CG, Boukes GJ, Cannon MD, Llambi F, Ryan D, et al. Systems analysis of BCL2 protein family interactions establishes a model to predict responses to chemotherapy. Cancer Res 2013; 73: 519–28.

17. Bradley CA, Dunne PD, Bingham V, McQuaid S, Khawaja H, Craig S, et al. Transcriptional upregulation of c-MET is associated with invasion and tumor budding in colorectal cancer. Oncotarget 2016; 7: 78932–45.

18. Chou TC,Talalay P. Quantitative analysis of dose-effect relationships: the combined effects of multiple drugs or enzyme inhibitors. Adv Enzyme Regul 1984; 22: 27–55.

19. Romanelli S, Perego P, Pratesi G, Carenini N, Tortoreto M,Zunino F. In vitro and in vivo interaction between cisplatin and topotecan in ovarian carcinoma systems. Cancer chemotherapy and pharmacology 1998; 41: 385–90.

20. Medico E, Russo M, Picco G, Cancelliere C, Valtorta E, Corti G, et al. The molecular landscape of colorectal cancer cell lines unveils clinically actionable kinase targets. Nature communications 2015; 6: 7002.

21. Mouradov D, Sloggett C, Jorissen RN, Love CG, Li S, Burgess AW, et al. Colorectal cancer cell lines are representative models of the main molecular subtypes of primary cancer. Cancer Res 2014; 74: 3238–47.

22. Ku JL, Shin YK, Kim DW, Kim KH, Choi JS, Hong SH, et al. Establishment and characterization of 13 human colorectal carcinoma cell lines: mutations of genes and expressions of drug-sensitivity genes and cancer stem cell markers. Carcinogenesis 2010; 31: 1003–9.

23. Liu Y,Bodmer WF. Analysis of P53 mutations and their expression in 56 colorectal cancer cell lines. Proc Natl Acad Sci U S A 2006; 103: 976–81.

24. Guinney J, Dienstmann R, Wang X, de Reynies A, Schlicker A, Soneson C, et al. The consensus molecular subtypes of colorectal cancer. Nat Med 2015; 21: 1350–6.

25. Zeng M, Lu J, Li L, Feru F, Quan C, Gero TW, et al. Potent and Selective Covalent Quinazoline Inhibitors of KRAS G12C. Cell Chem Biol 2017; 24: 1005–16 e3.

26. Rixe O,Fojo T. Is cell death a critical end point for anticancer therapies or is cytostasis sufficient? Clin Cancer Res 2007; 13: 7280–7.

27. Lindner AU, Salvucci M, Morgan C, Monsefi N, Resler AJ, Cremona M, et al. BCL-2 system analysis identifies high-risk colorectal cancer patients. Gut 2017; 66: 2141–8.

28. Awad MM, Liu S, Rybkin, II, Arbour KC, Dilly J, Zhu VW, et al. Acquired Resistance to KRAS(G12C) Inhibition in Cancer. N Engl J Med 2021; 384: 2382–93.

29. Tanaka N, Lin JJ, Li C, Ryan MB, Zhang J, Kiedrowski LA, et al. Clinical Acquired Resistance to KRAS(G12C) Inhibition through a Novel KRAS Switch-II Pocket Mutation and Polyclonal Alterations Converging on RAS-MAPK Reactivation. Cancer discovery 2021; 11: 1913–22.

30. Van Schaeybroeck S, Kyula JN, Fenton A, Fenning CS, Sasazuki T, Shirasawa S, et al. Oncogenic Kras promotes chemotherapy-induced growth factor shedding via ADAM17. Cancer Res 2011; 71: 1071–80.

31. Kyula JN, Van Schaeybroeck S, Doherty J, Fenning CS, Longley DB,Johnston PG. Chemotherapy-Induced Activation of ADAM-17: A Novel Mechanism of Drug Resistance in Colorectal Cancer. Clin Cancer Res 2010; 16: 3378–89.

32. Hallin J, Engstrom LD, Hargis L, Calinisan A, Aranda R, Briere DM, et al. The KRAS(G12C) Inhibitor MRTX849 Provides Insight toward Therapeutic Susceptibility of KRAS-Mutant Cancers in Mouse Models and Patients. Cancer discovery 2020; 10: 54–71.

33. Canon J, Rex K, Saiki AY, Mohr C, Cooke K, Bagal D, et al. The clinical KRAS(G12C) inhibitor AMG 510 drives anti-tumour immunity. Nature 2019; 575: 217–23.

34. Jänne PARI, Spira AI, Riely GJ, Papadopoulos KP, Sabari JK,. KRYSTAL-1: Activity and Safety of Adagrasib (MRTX849) in Advanced/ Metastatic Non–Small-Cell Lung Cancer (NSCLC) Harboring KRAS G12C Mutation. Eur J Cancer 2020 Oct;138:S1– 2.

35. Johnson ML OS, Barve M, Rybkin II, Papadopoulos KP, Leal TA. KRYSTAL-1: Activity and Safety of Adagrasib (MRTX849) in Patients with Colorectal Cancer (CRC) and Other Solid Tumors Harboring a KRAS G12C Mutation. Eur J Cancer 2020 Oct;138:S2.

36. Blair HA. Sotorasib: First Approval. Drugs 2021; 81: 1573–9.

37. Vaishnavi A, Scherzer MT, Kinsey CG, Parkman GL, Truong A, Ghazi P, et al. Inhibition of MEK1/2 Forestalls the Onset of Acquired Resistance to Entrectinib in Multiple Models of NTRK1-Driven Cancer. Cell reports 2020; 32: 107994.

38. Faber AC, Farago AF, Costa C, Dastur A, Gomez-Caraballo M, Robbins R, et al. Assessment of ABT-263 activity across a cancer cell line collection leads to a potent combination therapy for small-cell lung cancer. Proc Natl Acad Sci U S A 2015; 112: E1288–96.

39. Misale S, Fatherree JP, Cortez E, Li C, Bilton S, Timonina D, et al. KRAS G12C NSCLC Models Are Sensitive to Direct Targeting of KRAS in Combination with PI3K Inhibition. Clin Cancer Res 2019; 25: 796–807.

40. Faber AC, Corcoran RB, Ebi H, Sequist LV, Waltman BA, Chung E, et al. BIM expression in treatment-naive cancers predicts responsiveness to kinase inhibitors. Cancer discovery 2011; 1: 352–65.

41. Costa C, Molina MA, Drozdowskyj A, Gimenez-Capitan A, Bertran-Alamillo J, Karachaliou N, et al. The impact of EGFR T790M mutations and BIM mRNA expression on outcome in patients with EGFR-mutant NSCLC treated with erlotinib or chemotherapy in the randomized phase III EURTAC trial. Clin Cancer Res 2014; 20: 2001–10.

42. Ryan MB, Fece de la Cruz F, Phat S, Myers DT, Wong E, Shahzade HA, et al. Vertical Pathway Inhibition Overcomes Adaptive Feedback Resistance to KRAS(G12C) Inhibition. Clin Cancer Res 2020; 26: 1633–43.

43. Molina-Arcas M, Moore C, Rana S, van Maldegem F, Mugarza E, Romero-Clavijo P, et al. Development of combination therapies to maximize the impact of KRAS-G12C inhibitors in lung cancer. Science translational medicine 2019; 11.

44. Weber K, Harper N, Schwabe J,Cohen GM. BIM-mediated membrane insertion of the BAK pore domain is an essential requirement for apoptosis. Cell reports 2013; 5: 409–20.

45. Ramesh P, Lannagan TRM, Jackstadt R, Atencia Taboada L, Lansu N, Wirapati P, et al. BCL-XL is crucial for progression through the adenoma-to-carcinoma sequence of colorectal cancer. Cell Death Differ 2021; 28: 3282–96.

46. Corcoran RB, Cheng KA, Hata AN, Faber AC, Ebi H, Coffee EM, et al. Synthetic lethal interaction of combined BCL-XL and MEK inhibition promotes tumor regressions in KRAS mutant cancer models. Cancer Cell 2013; 23: 121–8.

47. Minn AJ, Rudin CM, Boise LH,Thompson CB. Expression of bcl-xL can confer a multidrug resistance phenotype. Blood 1995; 86: 1903–10.

48. Weiss J Yrd, Johnson M.L, et al. KRYSTAL-1: Adagrasib (MRTX849) as monotherapy or combined with cetuximab (Cetux) in patients (Pts) with colorectal cancer (CRC) harboring a KRASG12C mutation. Presented at: 2021 European Society for Medical Oncology Congress; September 16-21, 2021; virtual. Abstract LBA6.

49. Corcoran RB DK, Cleary JM, Parikh AR, Yeku OO, Weekes CD, et al. Phase I/II study of combined BCL-XL and MEK inhibition with navitoclax (N) and trametinib (T) in KRAS or NRAS mutant advanced solid tumours. Ann Oncol. 2019 Oct 1;30:v164.

50. Khan S, Wiegand J, Zhang P, Hu W, Thummuri D, Budamagunta V, et al. BCL-XL PROTAC degrader DT2216 synergizes with sotorasib in preclinical models of KRAS(G12C)-mutated cancers. J Hematol Oncol 2022; 15: 23.

