## Supplementary Figures for "Bcl-xL is a key mediator of apoptosis following KRAS^G12C^ inhibition in *KRAS*^*G12C*^ mutant colorectal cancer"

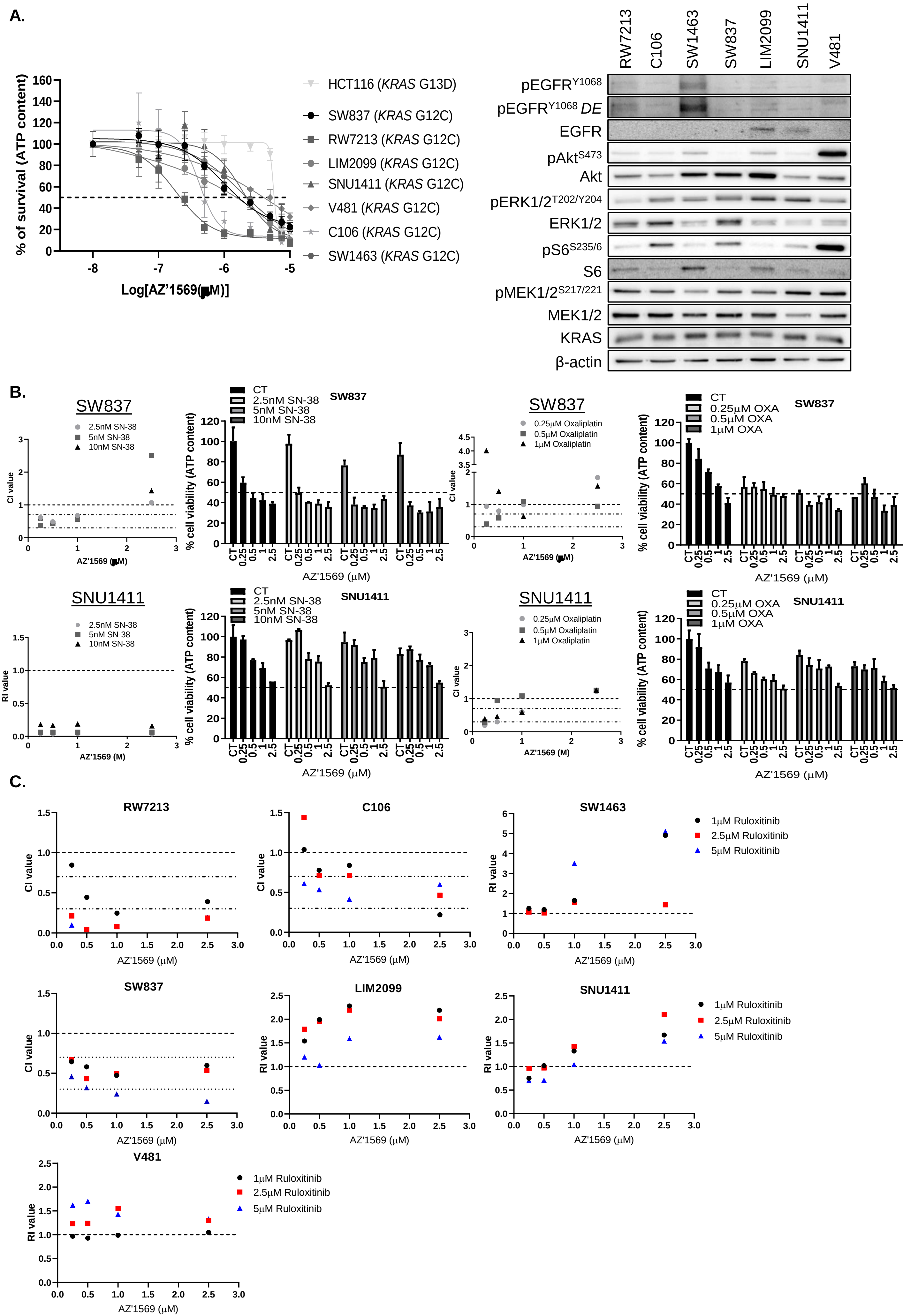
A.
% of survival (ATP content)
Log[AZ'1569 (μM)]
HCT116 (KRAS G13D)
SW837 (KRAS G12C)
RW7213 (KRAS G12C)
LIM2099 (KRAS G12C)
SNU1411 (KRAS G12C)
V481 (KRAS G12C)
C106 (KRAS G12C)
SW1463 (KRAS G12C)
RW7213
C106
SW1463
SW837
LIM2099
SNU1411
V481
pEGFR Y1068
pEGFR Y1068 DE
EGFR
pAkt S473
Akt
pERK1/2 T202/Y204
ERK1/2
pS6 S235/6
S6
pMEK1/2 S217/221
MEK1/2
KRAS
β-actin
B.
SW837
2.5nM SN-38
5nM SN-38
10nM SN-38
CI value
AZ'1569 (μM)
CT
% cell viability (ATP content)
0.25μM Oxaliplatin
0.5μM Oxaliplatin
1μM Oxaliplatin
0.25μM OXA
0.5μM OXA
1μM OXA
SNU1411
RI value
AZ'1569 (M)
C.
RW7213
C106
SW1463
SW837
LIM2099
SNU1411
V481
1μM Ruloxitinib
2.5μM Ruloxitinib
5μM Ruloxitinib
CI value
RI value
AZ'1569 (μM)

D.

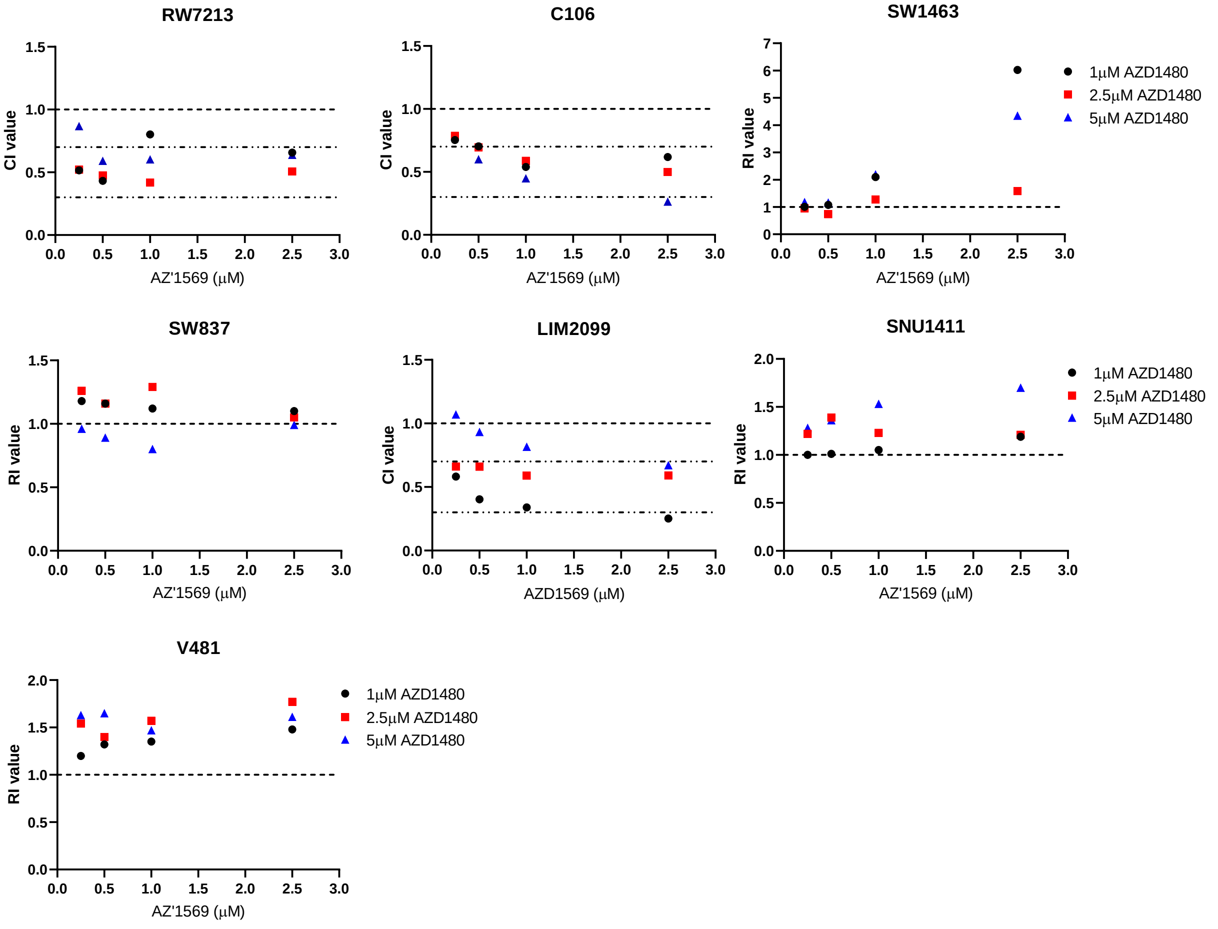

RW7213
CI value
AZ'1569 (μM)
C106
CI value
AZ'1569 (μM)
SW1463
RI value
AZ'1569 (μM)
1μM AZD1480
2.5μM AZD1480
5μM AZD1480
SW837
RI value
AZ'1569 (μM)
LIM2099
CI value
AZD1569 (μM)
SNU1411
RI value
AZ'1569 (μM)
1μM AZD1480
2.5μM AZD1480
5μM AZD1480
V481
RI value
AZ'1569 (μM)
1μM AZD1480
2.5μM AZD1480
5μM AZD1480


E.

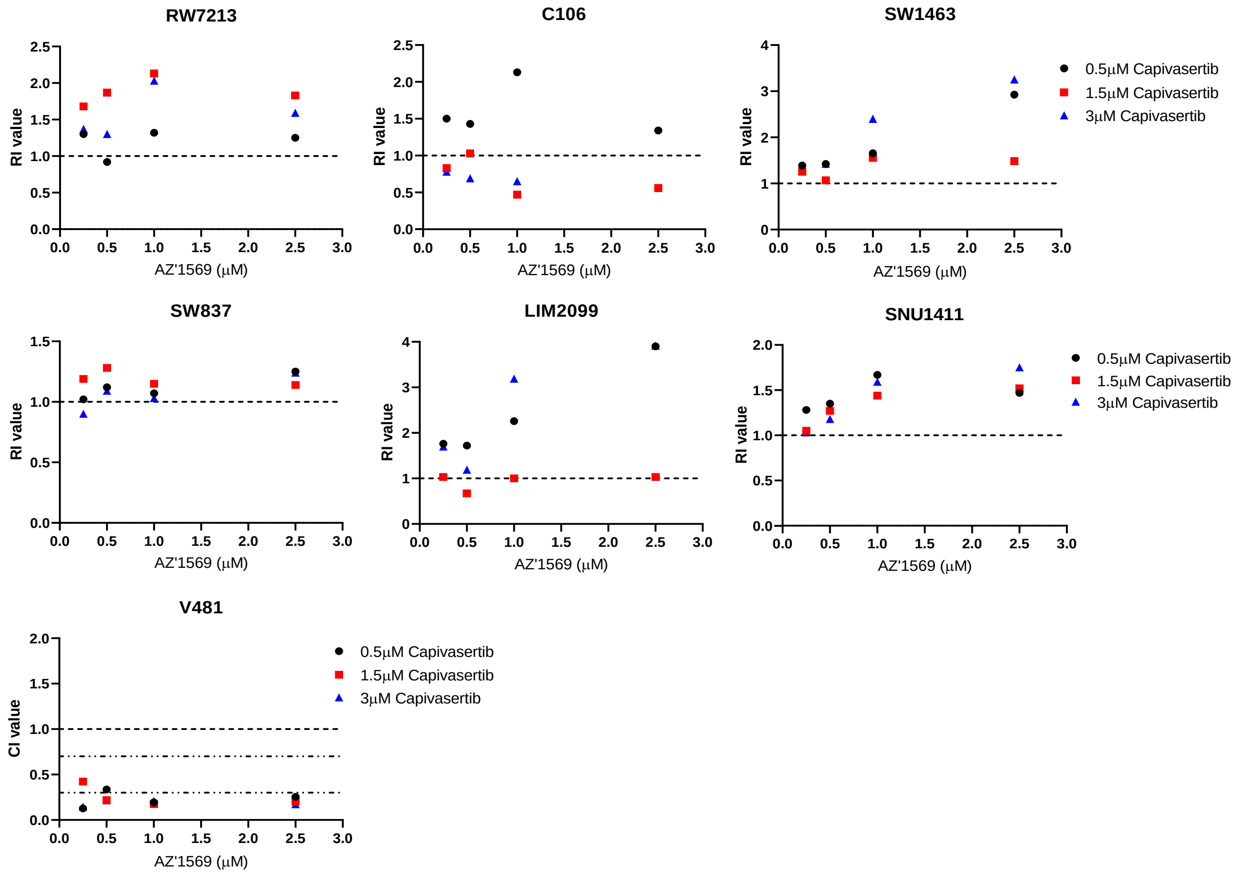

RW7213
RI value
AZ'1569 (μM)
C106
RI value
AZ'1569 (μM)
SW1463
RI value
AZ'1569 (μM)
0.5μM Capivasertib
1.5μM Capivasertib
3μM Capivasertib
SW837
RI value
AZ'1569 (μM)
LIM2099
RI value
AZ'1569 (μM)
SNU1411
RI value
AZ'1569 (μM)
0.5μM Capivasertib
1.5μM Capivasertib
3μM Capivasertib
V481
CI value
AZ'1569 (μM)
0.5μM Capivasertib
1.5μM Capivasertib
3μM Capivasertib

F.

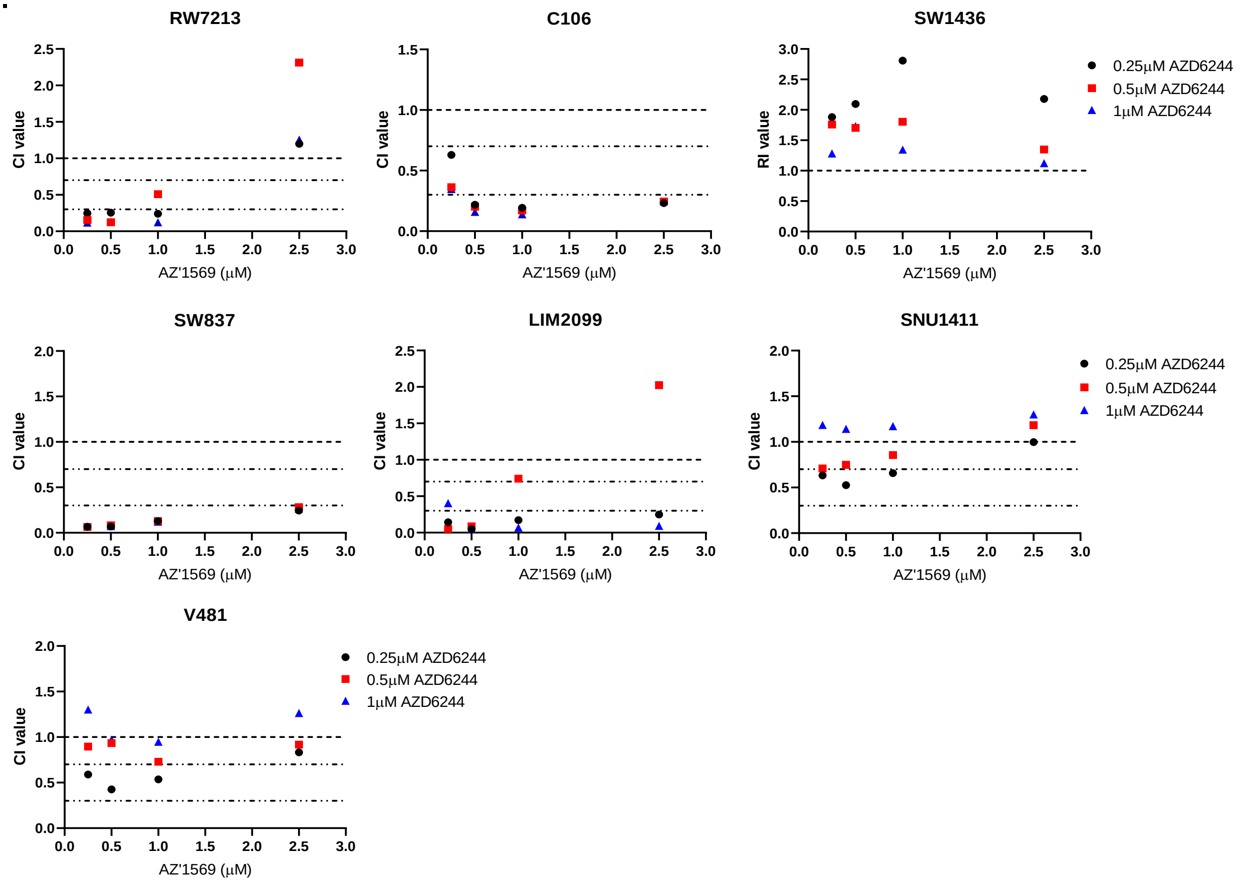
RW7213
C106
SW1436
SW837
LIM2099
SNU1411
V481
CI value
RI value
AZ'1569 (μM)
0.25μM AZD6244
0.5μM AZD6244
1μM AZD6244
0.0
0.5
1.0
1.5
2.0
2.5
3.0

G.

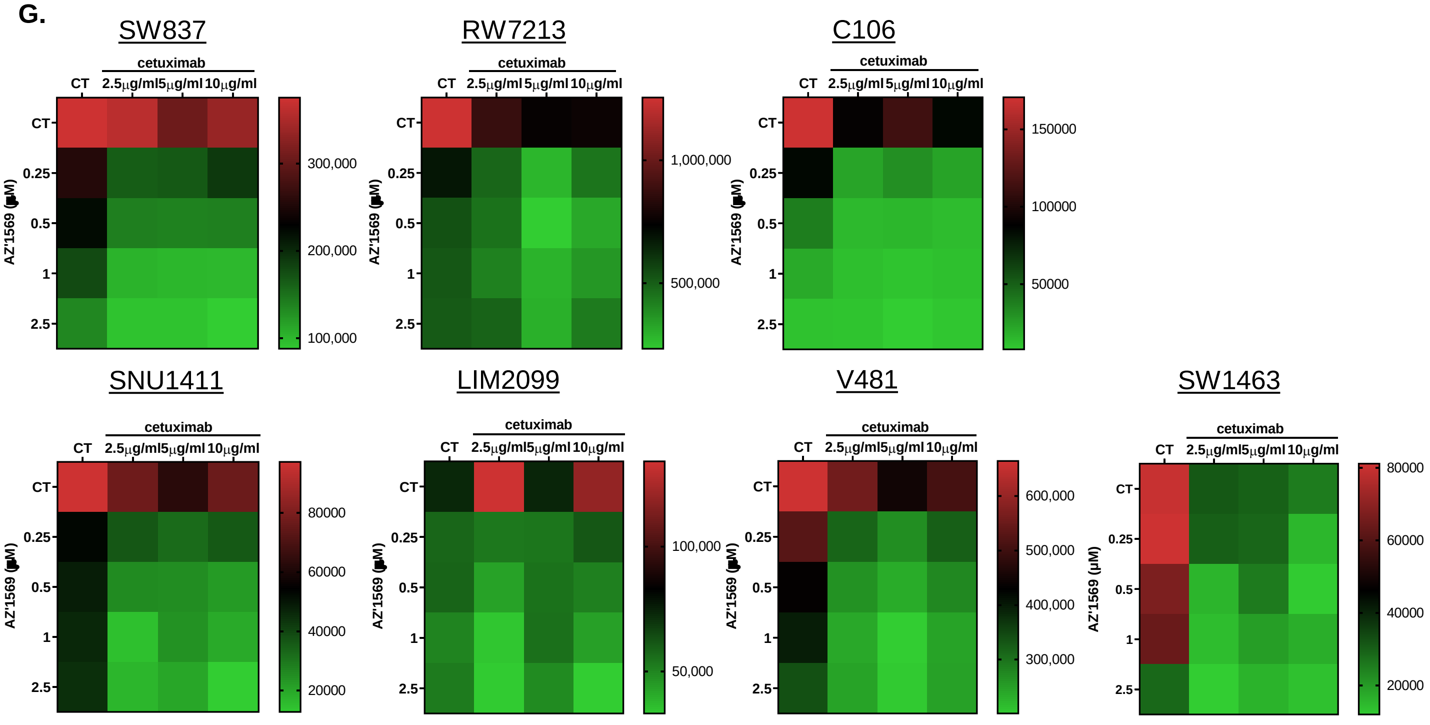
SW837
RW7213
C106
SNU1411
LIM2099
V481
SW1463
cetuximab
CT
2.5μg/ml
5μg/ml
10μg/ml
0.25
0.5
1
2.5
AZ'1569 (μM)
300,000
200,000
100,000
1,000,000
500,000
150000
100000
50000
80000
60000
40000
20000
100,000
50,000
600,000
500,000
400,000
300,000

**H.**

| Cell line | AZD6244 | AZD1480 | Ruloxitinib | Capivasertib | Cetuximab |
|---|---|---|---|---|---|
| **RW7213** | Moderate – high synergy | Slight-Moderate synergy | Moderate-high synergy | Additive – moderate synergy | High synergy |
| **C106** | Moderate – high synergy | Slight-Moderate synergy | Slight-Moderate synergy | Antagonistic - Additive | Moderate synergy |
| **SNU1411** | Additive – slight synergy | Additive – slight synergy | Additive – slight synergy | Additive – slight synergy | High synergy |
| **LIM2099** | Moderate – high synergy | Moderate synergy | Slight - Moderate synergy | Additive – moderate synergy | High synergy |
| **SW837** | High synergy | Additive | Moderate synergy | Additive | Moderate-high synergy |
| **SW1463** | Additive – moderate synergy | Additive – moderate synergy | Additive – moderate synergy | Additive – moderate synergy | Moderate synergy |
| **V481** | Additive - Slight synergy | Additive - Slight synergy | Additive - Slight synergy | High synergy | Moderate-high synergy |

**Supplementary Figure 1. Response of *KRAS*$^{G12C}$ MT CRC cells to KRAS$^{G12C}$ inhibitor AZ'1569 alone, combined with standard-of-care chemotherapies, cetuximab or inhibitors of the KRAS downstream effectors. A. Left:** *KRAS*$^{G12C}$ MT CRC cells were treated with increasing concentrations of AZ'1569 for 120h and cell viability determined using CellTiter-Glo® (CTG) assay. $IC_{50}$ values were calculated using Prism software package. Dashed line indicates 50% cell viability. Representative of three independent experiments is shown. **Right:** Basal expression levels of proteins within the EGFR/KRAS signalling axis, as determined by Western blotting (WB). DE=darker exposure. **B.** CTG assays in CRC cells treated with no drug (control), SN-38 or oxaliplatin, AZ'1569, SN-38 or oxaliplatin in combination with AZ'1569 for 72h. CI values were calculated using the method of Chou and Talalay. CI values >1, <1, and equal to 1 indicate antagonism, synergy and additive effects for drug combinations, respectively. Dashed lines indicate CI values of 0.3, 0.7 and 1. RI values were used where a compound had little/no effect on cell viability. RI values >1, <1, and equal to 1 indicate synergy, antagonism and additive effects for drug combinations, respectively. Absolute cell viability for different combinations is also shown. Dashed line indicates 50% cell viability. Representative results of at least three experiments are shown. **C, D, E, F, G**. *KRAS*$^{G12C}$ MT CRC cells were treated with AZ'1569 alone or combined with Ruloxitinib (**C**), AZD1480 (**D**), Capivasertib (**E**), AZD6244 (**F**) or cetuximab (**G**) for 120h and cell viability assessed using the CTG assay and CI/RI values were calculated. Heatmaps represent the absolute reduction in cell viability for combination of AZ'1569 with cetuximab. Representative results of at least three experiments are shown. **H.** Table summarising the nature of interaction between AZ'1569 and the different targeted drugs in the panel of *KRAS*$^{G12C}$ MT CRC cells.

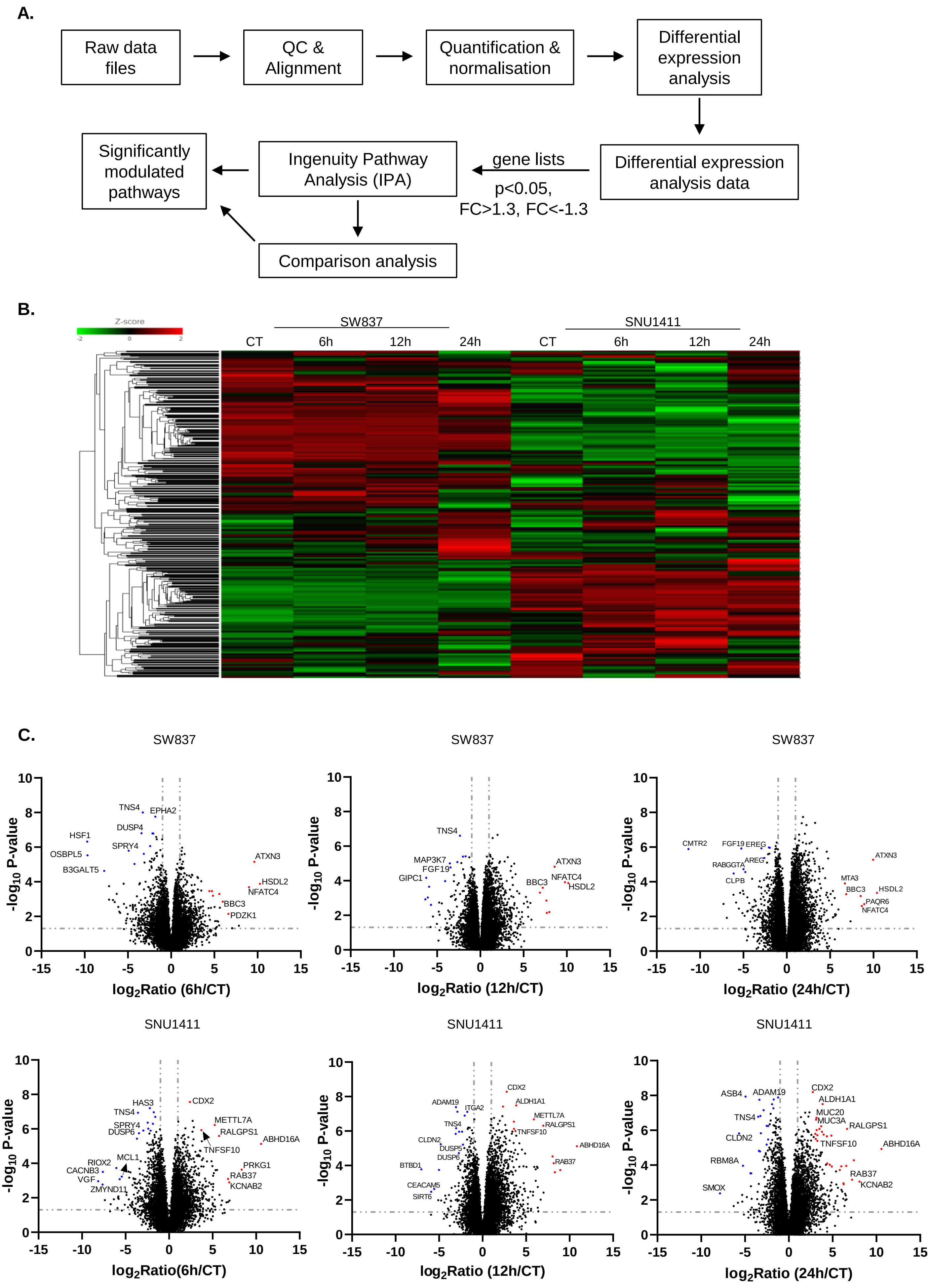

A.
Raw data files
QC & Alignment
Quantification & normalisation
Differential expression analysis
Differential expression analysis data
gene lists
p<0.05, FC>1.3, FC<-1.3
Ingenuity Pathway Analysis (IPA)
Significantly modulated pathways
Comparison analysis
B.
Z-score
-2
0
2
SW837
SNU1411
CT
6h
12h
24h
CT
6h
12h
24h
C.
SW837
SW837
SW837
SNU1411
SNU1411
SNU1411
-log10 P-value
log2Ratio (6h/CT)
log2Ratio (12h/CT)
log2Ratio (24h/CT)
log2Ratio(6h/CT)
TNS4
EPHA2
DUSP4
HSF1
SPRY4
OSBPL5
B3GALT5
ATXN3
HSDL2
NFATC4
BBC3
PDZK1
MAP3K7
FGF19
GIPC1
BBC3
NFATC4
HSDL2
CMTR2
FGF19
EREG
RABGGTA
AREG
CLPB
MTA3
BBC3
HSDL2
PAQR6
NFATC4
HAS3
TNS4
SPRY4
DUSP6
MCL1
RIOX2
CACNB3
VGF
ZMYND11
CDX2
METTL7A
RALGPS1
TNFSF10
ABHD16A
PRKG1
RAB37
KCNAB2
ADAM19
ITGA2
CLDN2
DUSP5
DUSP6
BTBD1
CEACAM5
SIRT6
ALDH1A1
ASB4
MUC20
MUC3A
RBM8A
SMOX

**D.**

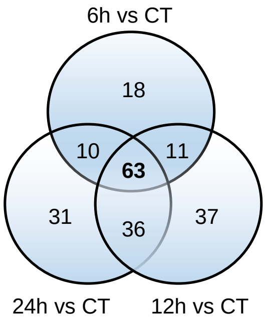


| | SW837 -$log_{10}$pvalue | | | SNU1411 -$log_{10}$pvalue | | |
|---|---|---|---|---|---|---|
| **Pathway** | **6h** | **12h** | **24h** | **6h** | **12h** | **24h** |
| Molecular Mechanisms of Cancer | 8.066559 | 9.160485 | 10.57428 | 10.04854 | 14.2449 | 6.75044 |
| EIF2 Signaling | 5.86876 | 7.00092 | 15.20365 | 6.62961 | 3.546702 | 1.897453 |
| Death Receptor Signaling | 8.000209 | 6.777285 | 4.362492 | 3.503648 | 4.676862 | 1.718967 |
| mTOR Signaling | 4.451432 | 3.765041 | 7.519563 | 5.400755 | 5.094452 | 4.531991 |
| Cell Cycle: G1/S Checkpoint Regulation | 5.620854 | 8.242366 | 7.647154 | 3.963368 | 5.37469 | 2.854804 |
| ERK/MAPK Signaling | 6.308891 | 3.00745 | 3.190258 | 3.053669 | 5.95694 | 7.065112 |
| Germ Cell-Sertoli Cell Junction Signaling | 4.627643 | 3.056515 | 3.808487 | 4.169748 | 5.01844 | 7.698286 |
| Sirtuin Signaling Pathway | 2.141829 | 6.681894 | 15.37388 | 6.373353 | 9.461571 | 12.54052 |
| HIPPO signaling | 4.517063 | 4.73105 | 6.743086 | 3.694088 | 3.779193 | 5.108474 |
| Glioblastoma Multiforme Signaling | 5.055339 | 2.185464 | 4.107408 | 3.106733 | 3.141882 | 2.303813 |
| Neuregulin Signaling | 4.347798 | 2.842146 | 2.174521 | 3.796415 | 4.082892 | 6.719828 |
| Chronic Myeloid Leukemia Signaling | 5.008688 | 4.347044 | 4.183806 | 2.848538 | 7.074057 | 4.845524 |
| Pancreatic Adenocarcinoma Signaling | 4.45597 | 3.411957 | 7.827764 | 3.377138 | 5.33906 | 2.276024 |
| HGF Signaling | 6.046239 | 3.243559 | 3.235769 | 1.773783 | 2.681906 | 3.605607 |
| Aryl Hydrocarbon Receptor Signaling | 3.425917 | 5.63355 | 4.986279 | 4.367558 | 6.098434 | 5.574729 |
| Cyclins and Cell Cycle Regulation | 4.461027 | 6.785328 | 6.56417 | 2.922904 | 6.117661 | 4.348443 |
| Senescence Pathway | 3.877711 | 7.316643 | 5.66488 | 3.343923 | 11.18359 | 9.807846 |
| Regulation of eIF4 and p70S6K Signaling | 3.53107 | 6.509098 | 9.571548 | 3.628492 | 6.061464 | 4.833019 |
| p53 Signaling | 4.166847 | 5.333847 | 4.889511 | 2.580736 | 3.86517 | 5.496942 |
| Integrin Signaling | 3.976514 | 1.6583 | 1.906902 | 2.732539 | 4.853825 | 10.56651 |
| PI3K/AKT Signaling | 2.841681 | 2.816287 | 2.321825 | 3.596005 | 6.352229 | 5.763325 |
| AMPK Signaling | 2.145715 | 5.839138 | 4.375414 | 4.210971 | 7.001451 | 6.317038 |
| Apoptosis Signaling | 4.079121 | 2.605405 | 1.922188 | 2.210349 | 2.754154 | 2.480502 |
| Necroptosis Signaling Pathway | 2.924407 | 3.725963 | 4.962115 | 3.332631 | 4.994035 | 3.056993 |
| Mouse Embryonic Stem Cell Pluripotency | 3.363275 | 1.544675 | 3.186304 | 2.848538 | 3.037925 | 3.050145 |
| Glioma Signaling | 4.36974 | 2.405081 | 3.044134 | 1.82709 | 4.817612 | 3.055761 |
| Hepatic Fibrosis Signaling Pathway | 3.32774 | 2.692935 | 2.223829 | 2.835887 | 3.463347 | 2.884307 |
| Huntington's Disease Signaling | 3.852445 | 4.171777 | 3.696712 | 2.098702 | 3.233941 | 2.990968 |
| RAR Activation | 1.810508 | 3.19748 | 3.113958 | 4.080867 | 8.03045 | 4.375325 |
| Superpathway of Inositol Phosphate Compounds | 2.316563 | 1.369722 | 3.263612 | 3.547439 | 7.212149 | 4.925232 |
| Telomerase Signaling | 3.81739 | 3.246768 | 2.498456 | 1.995155 | 3.663634 | 4.375888 |
| PDGF Signaling | 3.545288 | 1.52025 | 1.768099 | 2.193894 | 3.295954 | 2.074923 |
| Colorectal Cancer Metastasis Signaling | 3.107716 | 1.5551 | 2.278083 | 2.625719 | 3.440302 | 2.193115 |
| Protein Kinase A Signaling | 2.718927 | 5.403933 | 3.54176 | 2.984975 | 5.001706 | 6.027761 |
| Epithelial Adherens Junction Signaling | 3.516023 | 4.111612 | 2.915876 | 2.060211 | 3.542839 | 8.081199 |
| Wnt/β-catenin Signaling | 2.670487 | 2.009373 | 2.252591 | 2.900464 | 3.116873 | 2.504296 |
| Adipogenesis pathway | 2.41708 | 2.746657 | 3.500025 | 3.110097 | 4.480369 | 4.388986 |
| Regulation of the Epithelial-Mesenchymal Transition Pathway | 2.823253 | 2.157012 | 2.408112 | 2.620357 | 2.892655 | 1.729734 |
| ErbB2-ErbB3 Signaling | 2.865778 | 2.829327 | 1.533991 | 2.507073 | 2.93823 | 2.349024 |
| Estrogen Receptor Signaling | 2.481793 | 4.080669 | 3.6723 | 2.887593 | 6.565692 | 4.545054 |
| PTEN Signaling | 1.993212 | 2.451554 | 2.627861 | 3.374309 | 4.942862 | 5.239909 |
| FAK Signaling | 2.847932 | 1.749024 | 2.006656 | 2.484043 | 3.452761 | 4.625694 |
| ILK Signaling | 3.768393 | 2.468829 | 3.672179 | 1.48336 | 4.337454 | 3.584841 |
| ErbB Signaling | 3.291081 | 1.558632 | 1.614358 | 1.960085 | 2.557678 | 3.24187 |
| Hereditary Breast Cancer Signaling | 2.973333 | 4.460687 | 7.270296 | 2.198599 | 4.932814 | 5.170671 |
| ERK5 Signaling | 3.093571 | 4.124698 | 1.223689 | 1.645719 | 5.567041 | 5.205068 |
| Role of CHK Proteins in Cell Cycle Checkpoint Control | 3.177071 | 3.77546 | 7.299524 | 1.451807 | 6.128736 | 2.466845 |
| 14-3-3-mediated Signaling | 3.120791 | 2.944511 | 2.542891 | 1.462586 | 1.873977 | 3.737028 |
| Renal Cell Carcinoma Signaling | 2.839561 | 1.621615 | 2.317974 | 1.710091 | 2.836165 | 4.482347 |
| Cell Cycle Regulation by BTG Family Proteins | 2.400441 | 4.973676 | 4.107646 | 2.070882 | 5.593748 | 2.497677 |
| RhoA Signaling | 1.39444 | 2.375151 | 3.172541 | 2.965708 | 2.907493 | 7.296667 |
| Endometrial Cancer Signaling | 2.465495 | 3.388967 | 1.452742 | 1.867319 | 3.12552 | 2.890514 |
| Sertoli Cell-Sertoli Cell Junction Signaling | 2.382306 | 4.05599 | 5.261775 | 1.859315 | 5.705946 | 7.211127 |
| B Cell Receptor Signaling | 2.382306 | 3.231215 | 1.727015 | 1.666659 | 3.085727 | 1.355699 |
| PFKFB4 Signaling Pathway | 1.607917 | 2.893836 | 2.023514 | 2.382936 | 4.261462 | 1.817483 |
| IL-15 Production | 1.470977 | 2.787775 | 3.676793 | 2.238406 | 3.06506 | 1.749397 |
| Tight Junction Signaling | 1.905518 | 2.255402 | 2.383873 | 1.765485 | 6.736873 | 4.465814 |
| Non-Small Cell Lung Cancer Signaling | 2.243175 | 2.440347 | 1.611819 | 1.31747 | 3.19904 | 2.257231 |
| Role of BRCA1 in DNA Damage Response | 1.805638 | 1.368366 | 7.307542 | 1.710091 | 3.969201 | 1.974017 |
| Axonal Guidance Signaling | 1.772682 | 2.089048 | 2.595897 | 1.635474 | 5.259315 | 4.79811 |
| Role of PKR in Interferon Induction and Antiviral Response | 1.889369 | 3.403232 | 3.201751 | 1.480343 | 3.401816 | 2.235164 |
| Inhibition of ARE-Mediated mRNA Degradation Pathway | 1.43228 | 3.011667 | 2.467691 | 1.927513 | 5.004266 | 3.244749 |
| Unfolded protein response | 1.315897 | 2.586719 | 3.28722 | 1.843496 | 1.420228 | 2.578968 |
| Pyridoxal 5'-phosphate Salvage Pathway | 1.426833 | 2.454636 | 3.934055 | 1.517237 | 1.602019 | 2.703549 |

**E.**

| Pathway | Genes |
|---|---|
| Death receptor signalling | *ACIN1, ACTB, ACTG1, APAF1, ARHGDIB, BID, BIRC2, CASP10, CASP3, CASP6, CASP7, CASP8, CFLAR, DAXX, DFFB, HSPB1, IKBKG, LIMK1, MAP3K5, MAP4K4, MAPK8, NAIP, NFKBIA, NFKBID, PARP1, PARP10, PARP12, PARP14, PARP2, PARP4, PARP6, PARP8, PARP9, SPTAN1, TIPARP, TNFRSF1A, TNFRSF1B, TNFRSF21, TNFRSF25, TNFSF15, XIAP* |
| Apoptosis Signalling | *ACIN1, APAF1, BAD, BAK1, BCL2L1, BCL2L10, BIRC2, BIRC6, CAPN5, CAPNS1, CASP10, CASP7, CASP8, CDK1, DFFB, ENDOG, HTRA2, IKBKB, IKBKG, KRAS, MAP2K7, MAP3K5, MAP4K4, MAPK3, MAPK8, MCL1, NFKBIA, NFKBIB, PARP1, PLCG1, PLCG2, PRKCA, PRKCQ, RALB, RAP2B, RASD2, RRAS, SPTAN1, TNFRSF1A,TNFRSF1B, TP53, XIAP* |
| Necroptosis signalling | *AXL,BIRC2,BIRC3,CAMK2D,CAPN1,CAPN2,CAPN5,CAPNS1,CASP10,CASP8,CFLAR,CHUK,CYBB,CYLD,DAPK1,DNM1L,FKBP1A,GLUL,IKBKB,IKBKG,IRF3,IRF9,JAK1,JMJD7-PLA2G4B, MAP3K7, MERTK, PAM16, PLA2G10, PLA2G12A, PLA2G4B, PPP3CA, PPP3CB, RBCK1, RBL1, SHARPIN, SLC25A3, SLC25A4, SLC25A6, TAB2, TIMM17B, TIMM8B, TNFRSF1A, TNFRSF1B,TNFSF10,TNIP1,TOMM40,TOMM40L,TOMM7,TOMM70, TP53,TRADD,TSPO,TYRO3,UBC,VDAC2,VDAC3* |
| p53 signalling | *AKT1,AKT2,APAF1,ATM,BCL2L1,BIRC5,BRCA1,CCNK,CDK2,CHEK1, CHEK2, COQ8A, CSNK1D, CTNNB1, DRAM1, E2F1, GADD45B, GNL3,HIF1A,HIPK2,KAT2B,MAPK14,MDM4,PIK3C3, PIK3CA, IK3CB,PIK3R1,PIK3R2,PMAIP1,PML,PRKDC,PTEN,SCO2, SERPINB5, ERPINE2,SFN,THBS1,TP53,TP53INP1,TP73,TRIM29* |

**Supplementary Figure 2. KRAS$^{G12C}$ inhibition rewires the signalling network of *KRAS*$^{G12C}$ MT CRC cells. A.** Overview of the bioinformatics analysis of *in vitro* data in SW837 and SNU1411 cells to comprehensively map adaptive signalling following AZ'1569 inhibition. Generated FASTQ files were analysed using a workflow on Partek® Flow software, v10.0. Post-alignment QC and quantification of aligned reads to an annotation model (Partek E/M, default settings; min reads=10) was performed. Differential expression analysis was performed using the GSA (gene specific analysis) tool. A cut-off threshold of fold-change >1.3 or <-1.3, and p-value<0.05 was applied to gene lists. The resulting gene lists were imported into Ingenuity Pathway Analysis (IPA) software (Qiagen, UK) to identify significantly enriched pathways for each time-point following AZ'1569 treatment in both cell lines. Comparison analysis in IPA was used to compare significantly enriched (-$\log_{10}$ p-value >1.3) pathways across both cell lines. **B**. Two-dimensional hierarchical clustering analysis of genes that were induced or repressed following treatment of *KRAS*$^{G12C}$ MT SW837 and SNU1411 cells with AZ'1569 for 6h, 12h and 24h compared to the parental cells. **C**. Volcano plots show the up- and downregulated genes following AZ'1569 treatment in SW837 and SNU1411 cells at the indicated time-points. Dashed lines on the x and y-axis indicate $\log_2$ratio of 1/-1, and –$\log_{10}$ p-value=1.3, respectively. **D. Left:** Venn diagram of the up- and down regulated pathways at each of the time points in both SW837 and SNU1411 cells. CT = control. **Right**: Results of the IPA pathway analysis. List of the common 63 pathways altered following AZ'1569 treatment in both SW837 and SNU1411 cells at each of the time points. **E.** Table representing the significantly enriched pathways with gene sets in cell death-related signalling pathways.

**A.**

| CatalogNumber | Gene Symbol | GeneId | Description |
|---|---|---|---|
| L-015375-00 | ATG3 | 64422 | ATG3 ON-TARGETplus SMARTpool - Human |
| L-004374-00 | ATG5 | 9474 | ATG5 ON-TARGETplus SMARTpool - Human |
| L-020112-00 | ATG7 | 10533 | ATG7 ON-TARGETplus SMARTpool - Human |
| L-003870-00 | BAD | 572 | BAD ON-TARGETplus SMARTpool - Human |
| L-003305-00 | BAK1 | 578 | BAK1 ON-TARGETplus SMARTpool - Human |
| L-003308-01 | BAX | 581 | BAX ON-TARGETplus SMARTpool - Human |
| L-004380-00 | BBC3 | 27113 | BBC3 ON-TARGETplus SMARTpool - Human |
| L-003306-00 | BCL2A1 | 597 | BCL2A1 ON-TARGETplus SMARTpool - Human |
| L-004383-00 | BCL2L11 | 10018 | BCL2L11 ON-TARGETplus SMARTpool - Human |
| L-003458-00 | BCL2L1 | 598 | BCL2L1 ON-TARGETplus SMARTpool - Human |
| L-004384-00 | BCL2L2 | 599 | BCL2L2 ON-TARGETplus SMARTpool - Human |
| L-003307-00 | BCL2 | 596 | BCL2 ON-TARGETplus SMARTpool - Human |
| L-010552-00 | BECN1 | 8678 | BECN1 ON-TARGETplus SMARTpool - Human |
| L-004387-00 | BID | 637 | BID ON-TARGETplus SMARTpool - Human |
| L-004388-00 | BIK | 638 | BIK ON-TARGETplus SMARTpool - Human |
| L-004390-00 | BIRC2 | 329 | BIRC2 ON-TARGETplus SMARTpool - Human |
| L-004099-00 | BIRC3 | 330 | BIRC3 ON-TARGETplus SMARTpool - Human |
| L-004393-00 | BMF | 90427 | BMF ON-TARGETplus SMARTpool - Human |
| L-004394-00 | BOK | 666 | BOK ON-TARGETplus SMARTpool - Human |
| L-004402-00 | CASP10 | 843 | CASP10 ON-TARGETplus SMARTpool - Human |
| L-004401-00 | CASP1 | 834 | CASP1 ON-TARGETplus SMARTpool - Human |
| L-003465-00 | CASP2 | 835 | CASP2 ON-TARGETplus SMARTpool - Human |
| L-004307-00 | CASP3 | 836 | CASP3 ON-TARGETplus SMARTpool - Human |
| L-004406-00 | CASP6 | 839 | CASP6 ON-TARGETplus SMARTpool - Human |
| L-004407-00 | CASP7 | 840 | CASP7 ON-TARGETplus SMARTpool - Human |
| L-003466-00 | CASP8 | 841 | CASP8 ON-TARGETplus SMARTpool - Human |
| L-003309-00 | CASP9 | 842 | CASP9 ON-TARGETplus SMARTpool - Human |
| L-003772-00 | CFLAR | 8837 | CFLAR ON-TARGETplus SMARTpool - Human |
| L-003800-00 | FADD | 8772 | FADD ON-TARGETplus SMARTpool - Human |
| L-003776-00 | FAS | 355 | FAS ON-TARGETplus SMARTpool - Human |
| L-008216-00 | HRK | 8739 | HRK ON-TARGETplus SMARTpool - Human |
| L-003585-00 | MAP3K9 | 4293 | MAP3K9 ON-TARGETplus SMARTpool - Human |
| L-004501-00 | MCL1 | 4170 | MCL1 ON-TARGETplus SMARTpool - Human |
| L-005275-00 | PMAIP1 | 5366 | PMAIP1 ON-TARGETplus SMARTpool - Human |
| L-003533-00 | RELA | 5970 | RELA ON-TARGETplus SMARTpool - Human |
| L-004445-00 | RIPK1 | 8737 | RIPK1 ON-TARGETplus SMARTpool - Human |
| L-003534-00 | RIPK3 | 11035 | RIPK3 ON-TARGETplus SMARTpool - Human |
| L-003544-00 | STAT3 | 6774 | STAT3 ON-TARGETplus SMARTpool - Human |
| L-008090-00 | TNFRSF10A | 8797 | TNFRSF10A ON-TARGETplus SMARTpool - Human |
| L-004448-00 | TNFRSF10B | 8795 | TNFRSF10B ON-TARGETplus SMARTpool - Human |
| L-003329-00 | TP53 | 7157 | TP53 ON-TARGETplus SMARTpool - Human |
| L-004098-00 | XIAP | 331 | XIAP ON-TARGETplus SMARTpool - Human |

**B.**

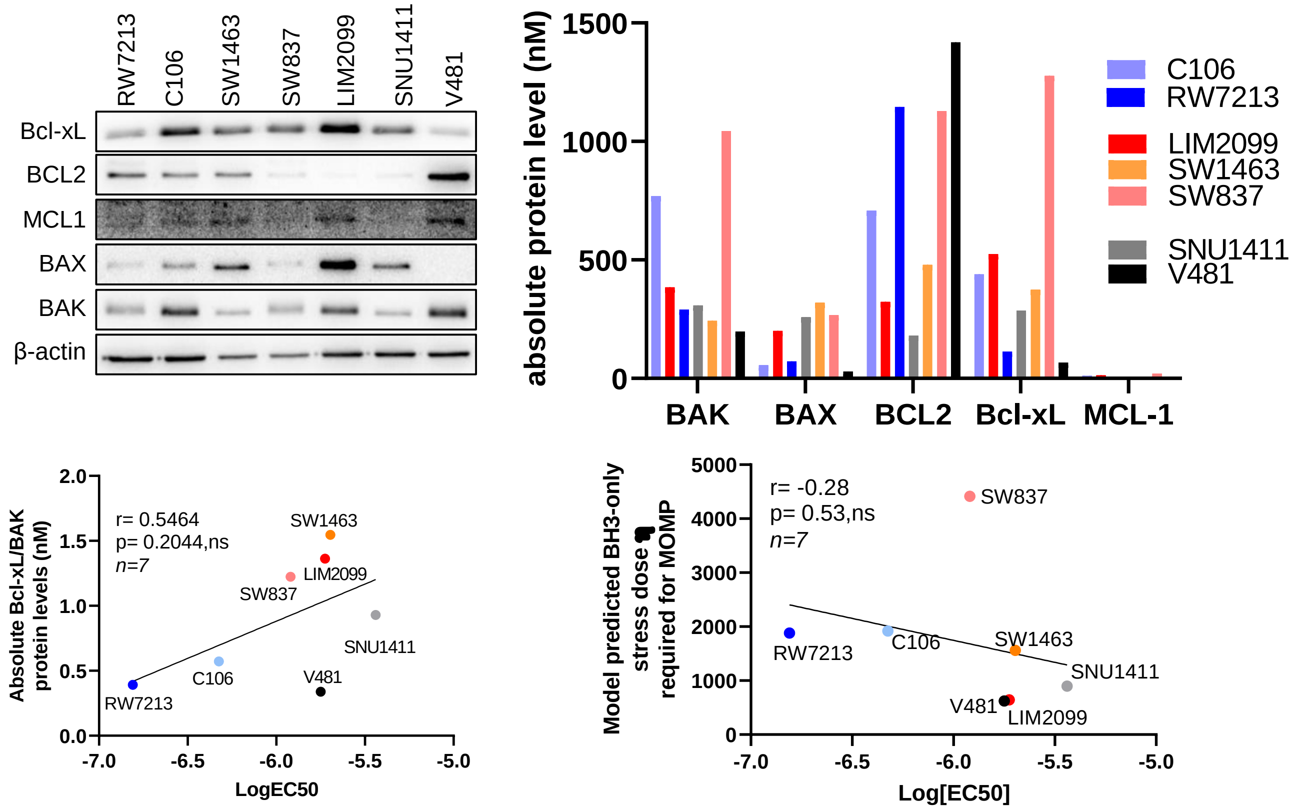

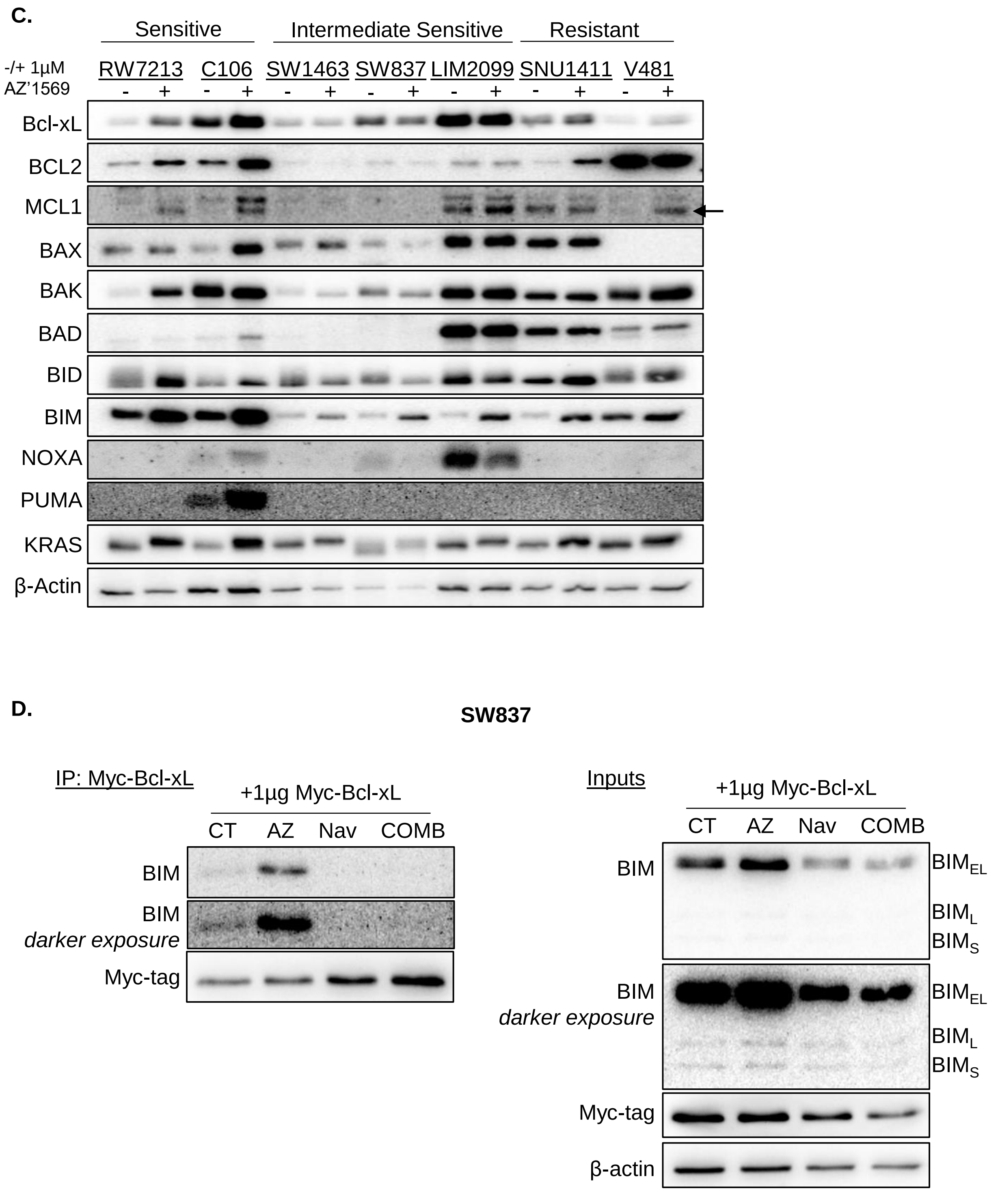


**Supplementary figure 3. Bcl-xL is a major escape pathway following KRAS$^{G12C}$ inhibition in *KRAS*$^{G12C}$ MT CRC cells. A.** Details of the ON-Targetplus siRNA library (Dharmacon) used to identify common targets of intrinsic resistance to KRAS$^{G12C}$ inhibition in SW837 and SNU1411 cells. **B. Top Left:** Basal expression levels of Bcl-xL, BCL2, MCL1, BAX and BAK, as determined by WB. **Top right**: Absolute protein levels of the BCL2 family proteins BAK, BAX, BCL2, Bcl-xL and MCL1. Protein levels obtained from quantitative WB were first normalised to β-actin and eventually normalised to HCT-116 cells to obtain absolute protein levels in nM. Protein levels in HCT-116 cells were previously quantified as described in Lindner *et al.* 2013 (Ref. 16). **Bottom left:** Pearson correlation of absolute protein Bcl-xL/BAK ratio plotted against Log[$EC_{50}$] AZ'1569 in the panel of *KRAS*$^{G12C}$ MT cell lines. **Bottom right:** Pearson correlation of the model predicted stress dose required to induce MOMP plotted against Log[$EC_{50}$] AZ'1569 in the panel of *KRAS*$^{G12C}$ MT CRC cell lines. **C. Left:** *KRAS*$^{G12C}$ MT cell lines were treated with 1µM AZ'1569 for 48h. Expression of BCL2 family members was determined by WB. **D.** SW837 cells were transfected with 1µg Myc-tagged Bcl-xL for 24 hours, followed by treatment with 1µM AZ'1569 (AZ), 1µM Navitoclax (Nav), or combination (COMB) for a further 24 hours. Cells were harvested and lysed and Myc-tagged Bcl-xL was immunoprecipitated. WB was used to determine the expression of Myc-tag, Bim and β-actin. Data is representative of three independent experiments.

A.

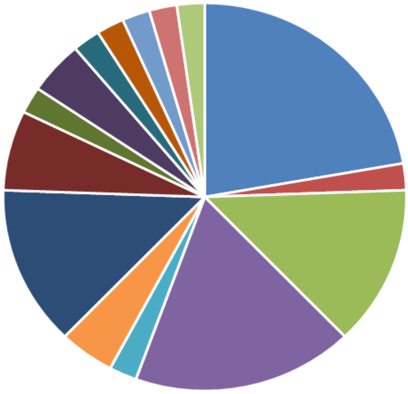


| Name | Pathway | Target/Process |
|---|---|---|
| ABT-737 | Apoptosis/cell death | Bcl-2, Bcl-xL, Bcl-w, ,Autophagy |
| Saracatinib (AZD0530) | Angiogenesis/ Src inhibitor | Src |
| Vorinostat (SAHA, MK0683) | Epigenetics | Autophagy,HDAC |
| Entinostat (MS-275) | Epigenetics | HDAC |
| Olaparib (AZD2281, Ku-0059436) | DNA damage | PARP |
| Nutlin-3 | MDM2 antagonist/ p53/apoptosis | E3 Ligase ,Mdm2 |
| Vismodegib (GDC-0449) | Wnt signalling pathway | Hedgehog/Smoothened |
| Alisertib (MLN8237) | Cell cycle | Aurora Kinase |
| Barasertib (AZD1152-HQPA) | Cell cycle | Aurora Kinase |
| Roscovitine (Seliciclib,CYC202) | Cell cycle | CDK |
| Ganetespib (STA-9090) | cytoskeletal signalling/ HSP90 inhibitor | HSP (e.g. HSP90) |
| BIBR 1532 | DNA damage | Telomerase |
| Epothilone A | cytoskeletal signalling | Microtubule Associated |
| AZD7762 | cell cycle | Chk |
| Ixazomib (MLN2238) | proteosome inhibitor | Proteasome |
| Degrasyn (WP1130) | proteosome inhibitor | Bcr-Abl,DUB |
| Rosiglitazone | Metabolism | PPAR |
| AT406 (SM-406) | Cell death/IAP | E3 Ligase ,IAP |
| I-BET151 (GSK1210151A) | Epigenetics | Epigenetic Reader Do |
| Sirtinol | Epigenetics | Sirtuin |
| Carfilzomib (PR-171) | proteosome inhibitor | Proteasome |
| IMD 0354 | NF-kB | IκB/IKK |
| Salubrinal | ER stress/ UPR | PERK |
| JNK-IN-8 | MAPK pathway | JNK |
| Birinapant | IAP inhibitor/cell death | IAP |
| RG-7112 | Apoptosis/Mdm2 inhibitor | Mdm2 |
| AZD1208 | JAK/STAT | Pim |
| UNC1999 | Epigenetics | Histone Methyltransferase |
| Tasisulam | caspase activator/cell death/ | Caspase |
| TH287 | DNA damage | MTH1 |
| AZD6738 | DNA damage | ATM/ATR |
| Venetoclax (ABT-199, GDC-0199) | Apoptosis | Bcl-2 |
| Sabutoclax | Apoptosis | Bcl-2, Bcl-xL, Mcl-1, Bfl-1 |
| CB-5083 | Transmembrane transporters | ATPase |
| AZD1390 | DNA damage | ATM/ATR |
| Palbociclib (PD-0332991) HCl | Cell cycle inhibitor | CDK |
| CX-5461 | DNA damage | DNA/RNA Synthesis |
| NU7026 | DNA damage | DNA-PK |
| Palifosfamide | DNA damage | DNA alkylator |
| EPZ5676 | Epigenetics | Histone Methyltransferase |
| AZD5991 | Cell death- Mcl-1 inhibitor | Mcl-1 |
| AZD4573 | Cell cycle | CDK9 inhibitor |
| Durvalumab | Immunology & Inflammation | PD-1/PDL-1 interaction |
| ONC206 | ER stress/ UPR | DRD2 anatagonist, ER stress inducer |
| iz-TRAIL | Cell death | death receptor agonist |

B.

| Drug - concentration (+1μM AZ'1569) | CI value SW837 | CI value SNU1411 |
|---|---|---|
| ONC206 - 1μM | 1.2 | 21.99 |
| Degrasyn - 2μM | 1.01 | 0.94 |
| RG-7112 - 1μM | 3.06 | 1.97 |
| CX-5461- 0.2μM | 0.69 | 1.53 |
| AT-406 - 1μM | 1.37* | 0.90 |
| IMD0354 - 0.5μM | 0.93 | 0.54 |
| AZD4573 - 1μM | 1.00 | 1.44 |
| CB-5083 - 1μM | 1.12 | 1.14 |
| UNC1999 - 3μM | 0.78 | 1.31 |
| JNK-IN-8 - 1μM | 0.56 | 1.44 |
| Entinostat - 1μM | 0.266 | 0.46 |
| ABT-737 - 1μM | 0.57 | 0.53 |

C.

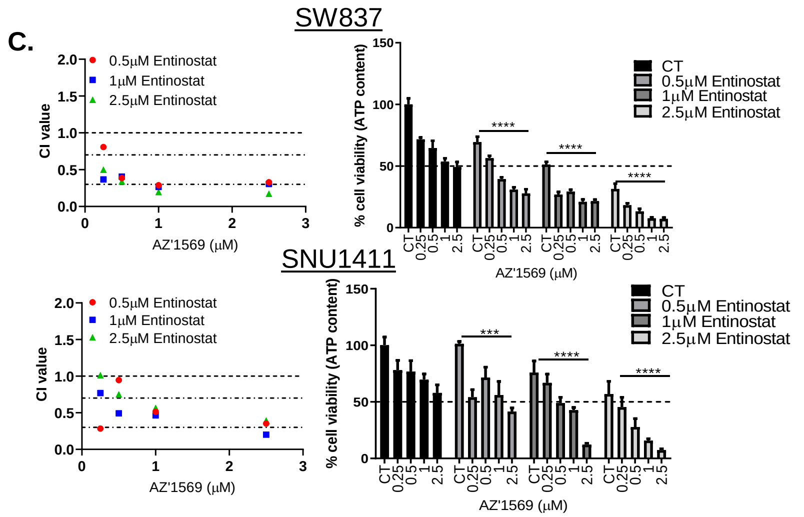


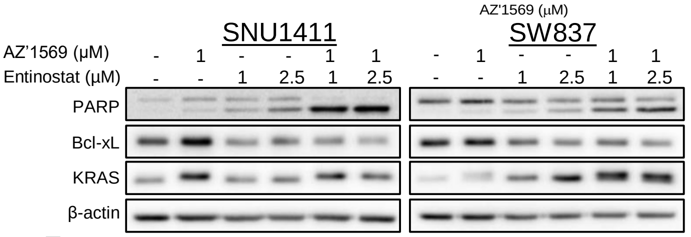


D.

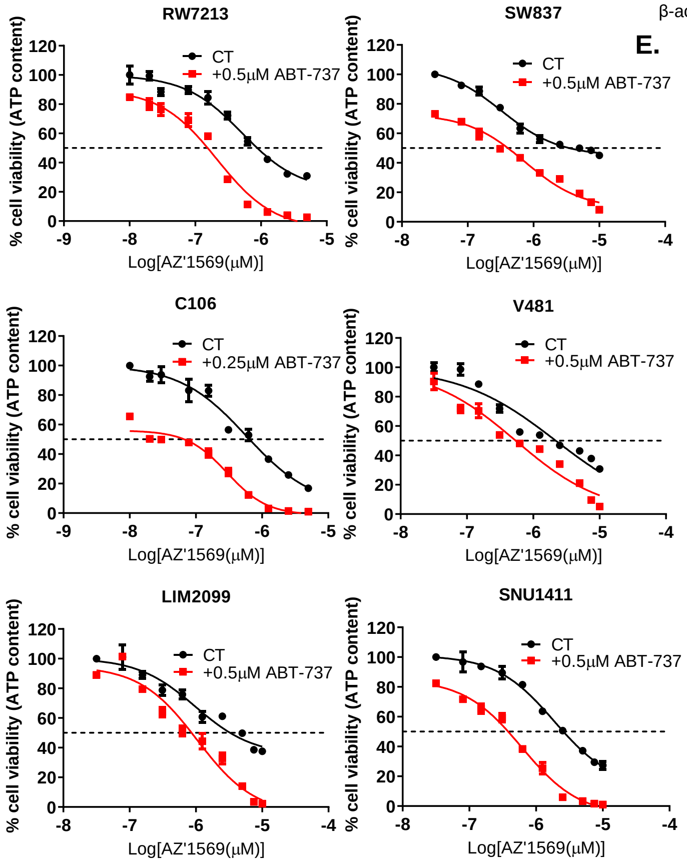


E.

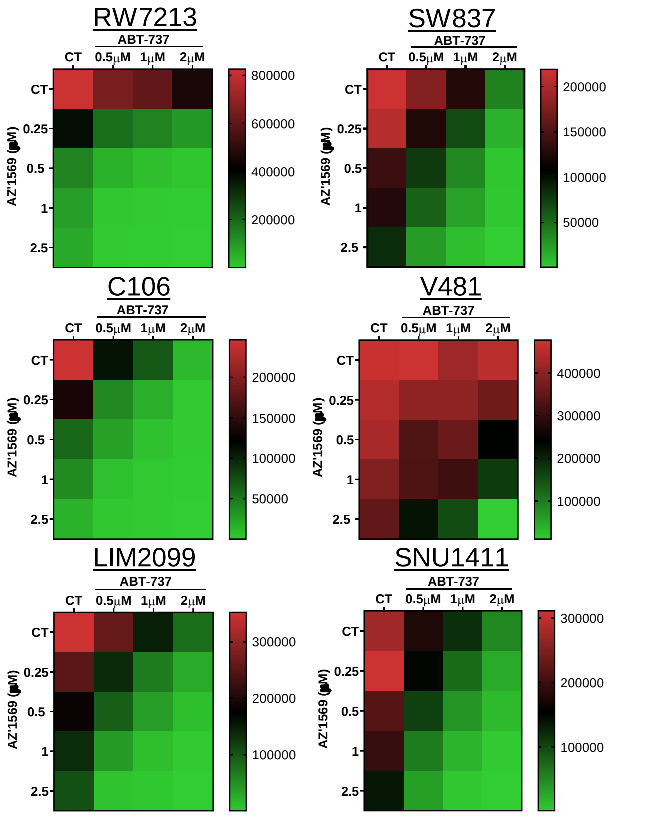


F.

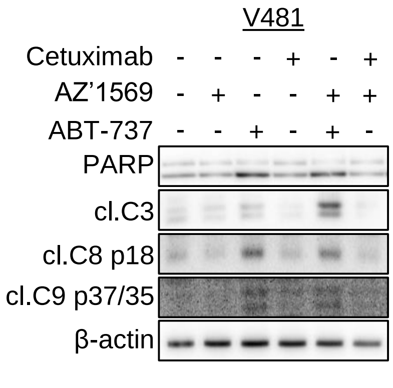


**Supplementary figure 4. High-throughput drug screening in *KRAS*$^{G12C}$ MT CRC cells. A. Top:** Pie chart of pathways targeted by drugs included in the compound screen. **Bottom:** Table with 45 drugs in the primary drug screen, and pathways/targets modulated by these compounds. **B**. Positive hits from primary screen. SNU1411 and SW837 cells were treated with AZ'1569 alone or combined with each of 12 compounds for 72h and CI/RI values determined. CI/RI values for one concentration of the 12 drugs with 1μM AZ'1569 is shown. * = RI value. **C. Upper/middle**: SW837 and SNU1411 were co-treated with Entinostat (drug 11) and AZ'1569 for 72h and CI values calculated. Absolute cell viability for different combinations is also shown. Dashed line indicates 50% cell viability. **Lower**: PARP, Bcl-xL and KRAS in CRC cells co-treated with Entinostat and AZ'1569. **D.** Dose response curves for AZ'1569 in absence and presence of ABT-737 (drug 12) for 72h. Cell viability was assessed using a CTG assay. Dashed lines represent 50% cell viability. **E.** CTG assays in CRC cells, co-treated with ABT-737 and AZ'1569 for 72h. Heatmaps show absolute cell viability. **F.** PARP, Cleaved-C3/C8/C9 in V481 cells treated with AZ'1569/cetuximab or AZ'1569/ABT-737. Data is representative of three independent experiments.

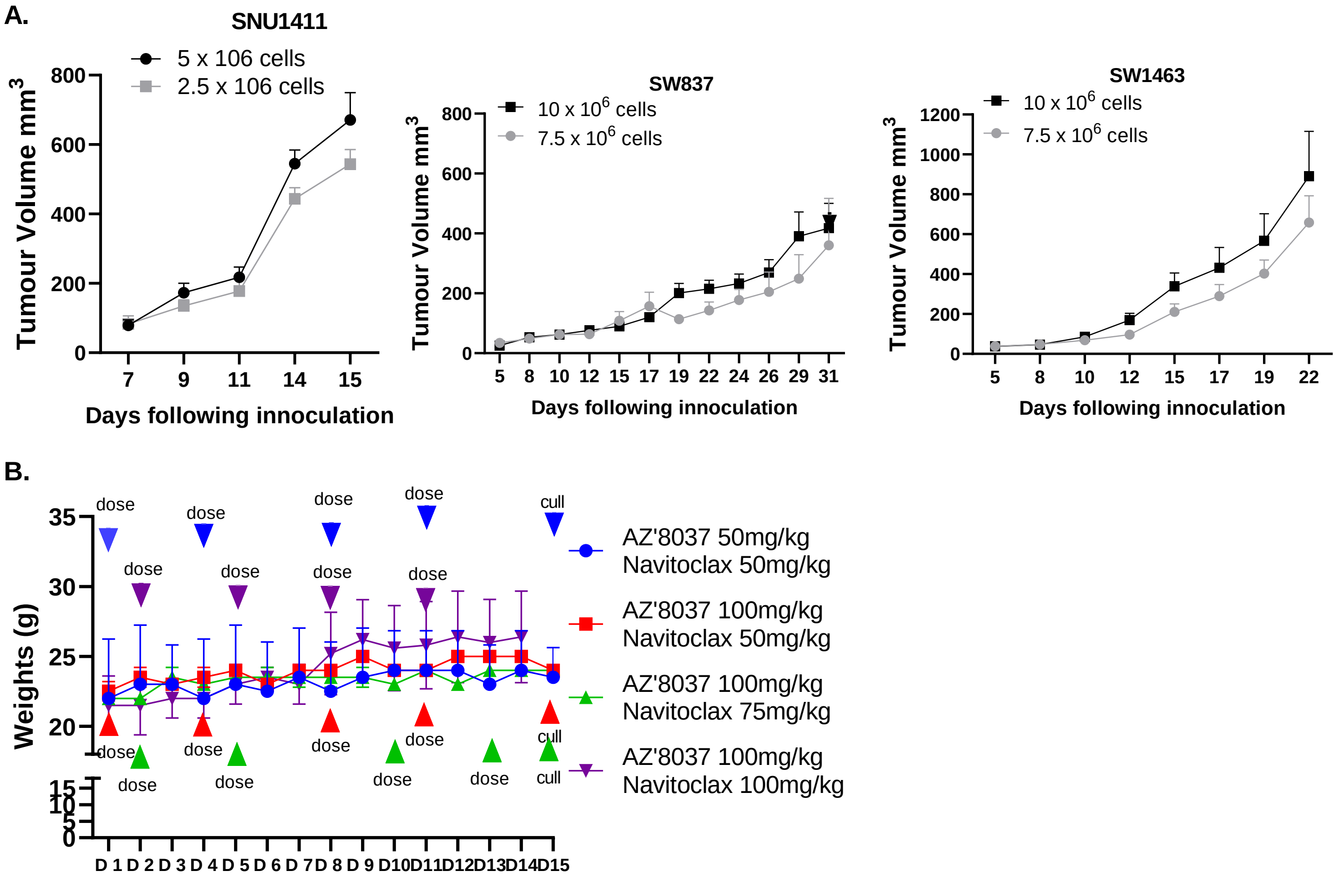


**Supplementary figure 5. Treatment of *KRAS*$^{G12C}$ MT CRC mouse models with AZ’8037, Navitoclax or combination. A.** Tumour growth curves in NOD-SCID mice for *KRAS*$^{G12C}$ MT SW837, SNU1411 and SW1463 CRC xenografts following inoculation with the indicated cell numbers. **B**. Weights of non-tumour-bearing NOD-SCID mice following 2 weeks treatment with AZ’8037/Navitoclax. Dose levels and days of treatments are indicated on the graphs.

**A.**

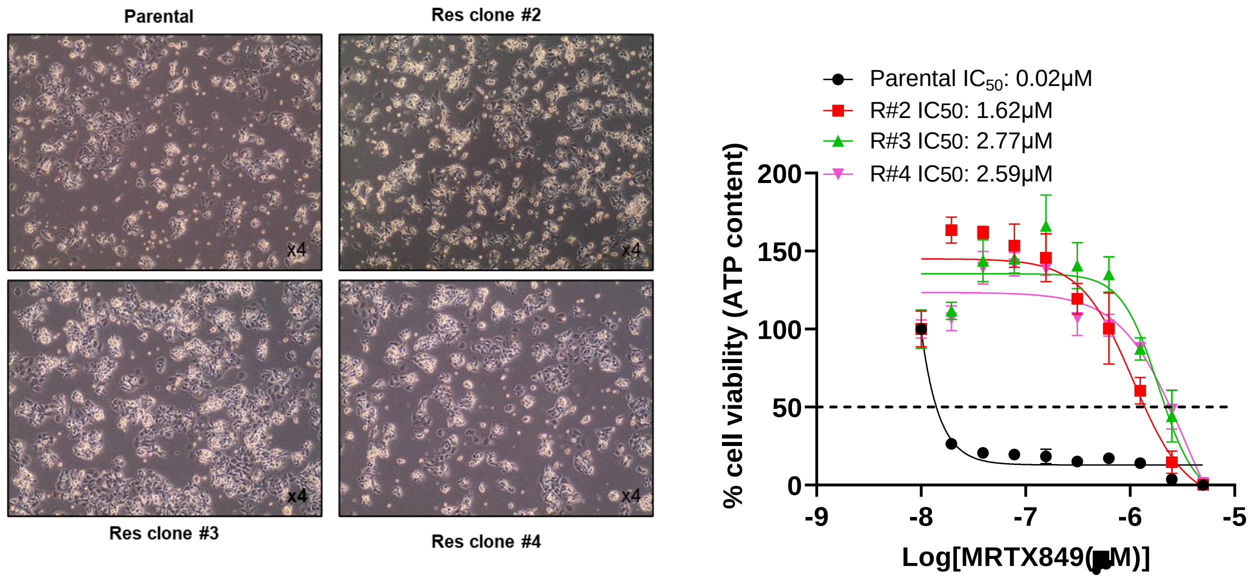

Parental
Res clone #2
Res clone #3
Res clone #4
x4
Parental IC50: 0.02μM
R#2 IC50: 1.62μM
R#3 IC50: 2.77μM
R#4 IC50: 2.59μM
% cell viability (ATP content)
200
150
100
50
0
-9
-8
-7
-6
-5
Log[MRTX849(μM)]


**B.**

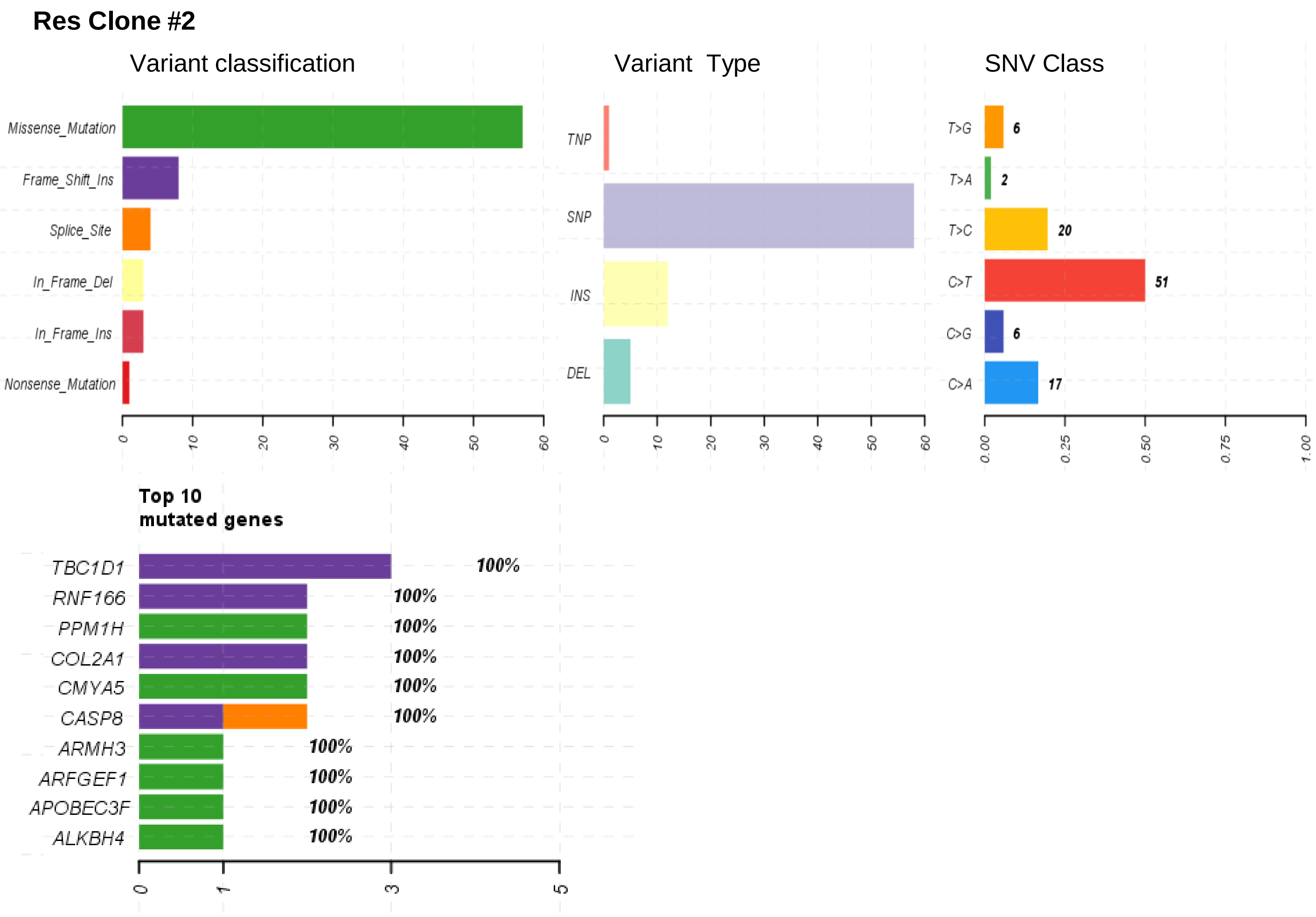

Res Clone #2
Variant classification
Missense_Mutation
Frame_Shift_Ins
Splice_Site
In_Frame_Del
In_Frame_Ins
Nonsense_Mutation
0
10
20
30
40
50
60
Variant Type
TNP
SNP
INS
DEL
SNV Class
T>G
6
T>A
2
T>C
20
C>T
51
C>G
6
C>A
17
0.00
0.25
0.50
0.75
1.00
Top 10 mutated genes
TBC1D1
100%
RNF166
100%
PPM1H
100%
COL2A1
100%
CMYA5
100%
CASP8
100%
ARMH3
100%
ARFGEF1
100%
APOBEC3F
100%
ALKBH4
100%
0
1
3
5

**B. continued**

### Res Clone #3

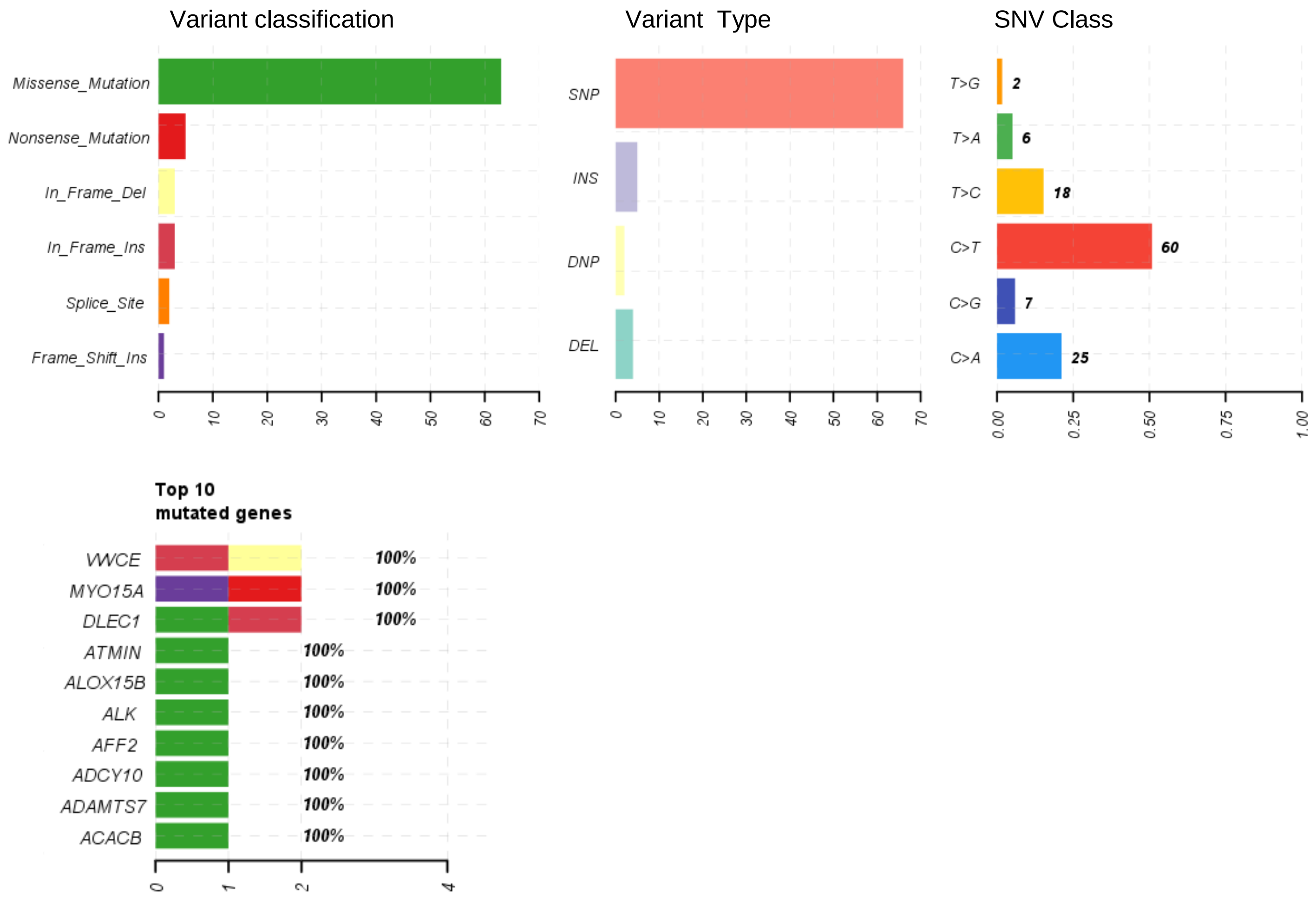


### Res Clone #4

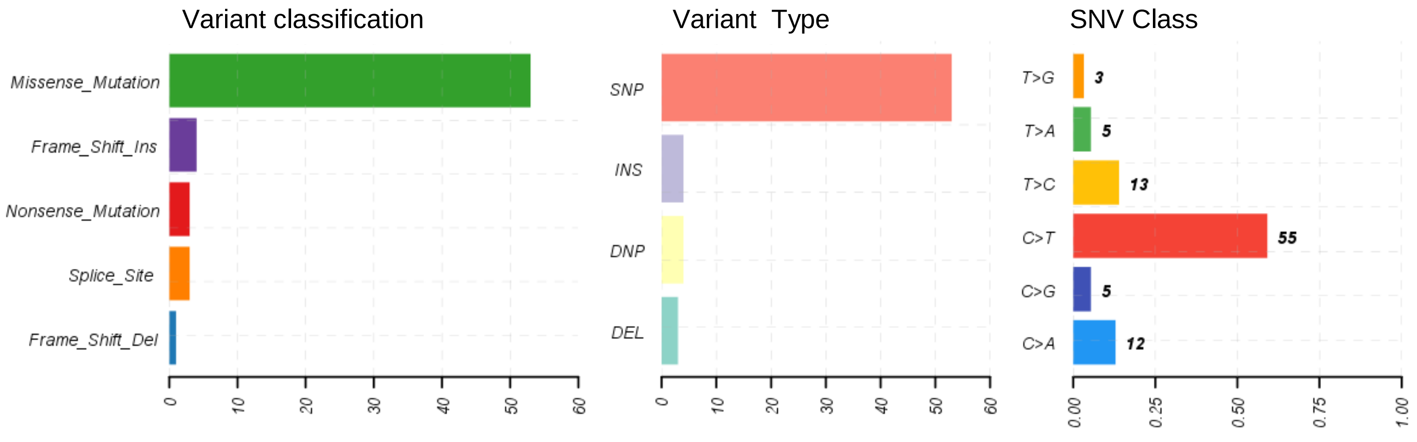


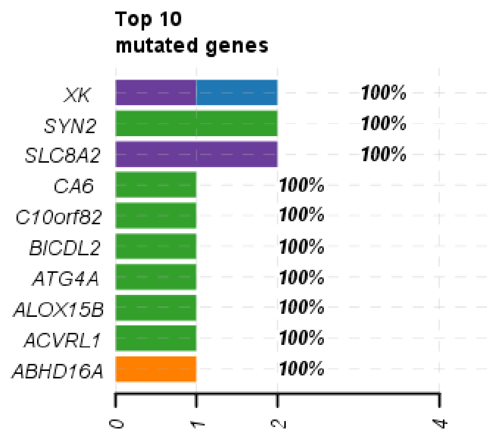


**C.**

| *KRAS*$^{G12C}$ CRC RW7213 | Additional basal gene alterations | Acquired mutations |
|---|---|---|
| Parental | *TP53* p.L257R, *BRCA2* amplification | None |
| Res clone #2 | *TP53* p.L257R, *BRCA2* amplification | *KRAS* amplification |
| Res clone #3 | *TP53* p.L257R, *BRCA2* amplification | *KRAS* amplification, *EGFR* amplification |
| Res clone #4 | *TP53* p.L257R, *BRCA2* amplification | *KRAS* amplification |

D.

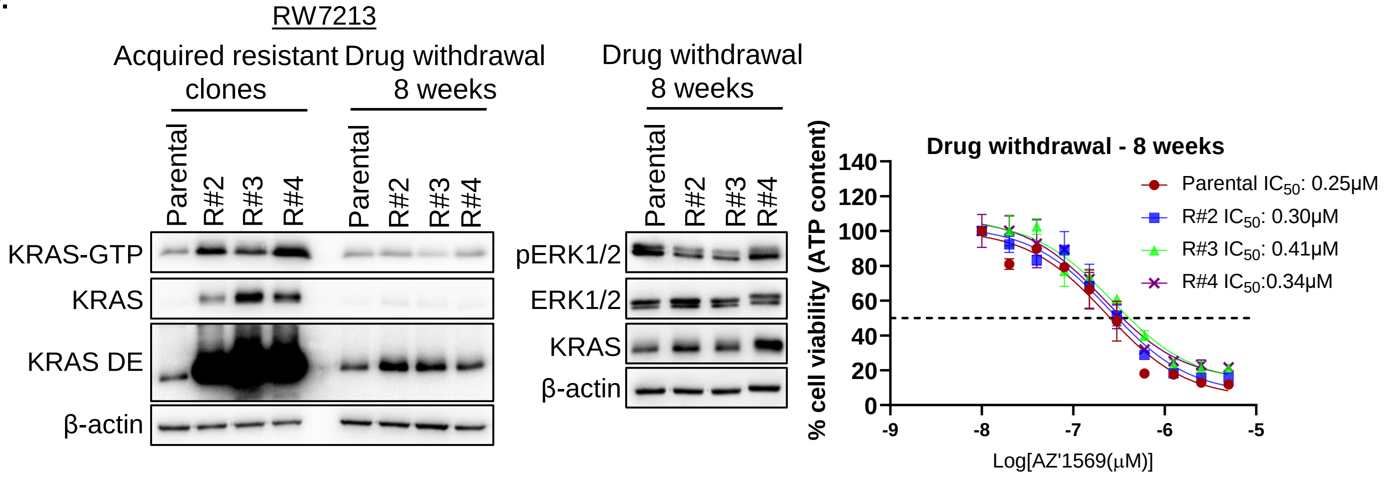
RW7213
Acquired resistant clones
Drug withdrawal 8 weeks
Drug withdrawal 8 weeks
Parental
R#2
R#3
R#4
KRAS-GTP
KRAS
KRAS DE
β-actin
pERK1/2
ERK1/2
KRAS
β-actin
Drug withdrawal - 8 weeks
Parental IC50: 0.25μM
R#2 IC50: 0.30μM
R#3 IC50: 0.41μM
R#4 IC50:0.34μM
% cell viability (ATP content)
Log[AZ'1569(μM)]

E.

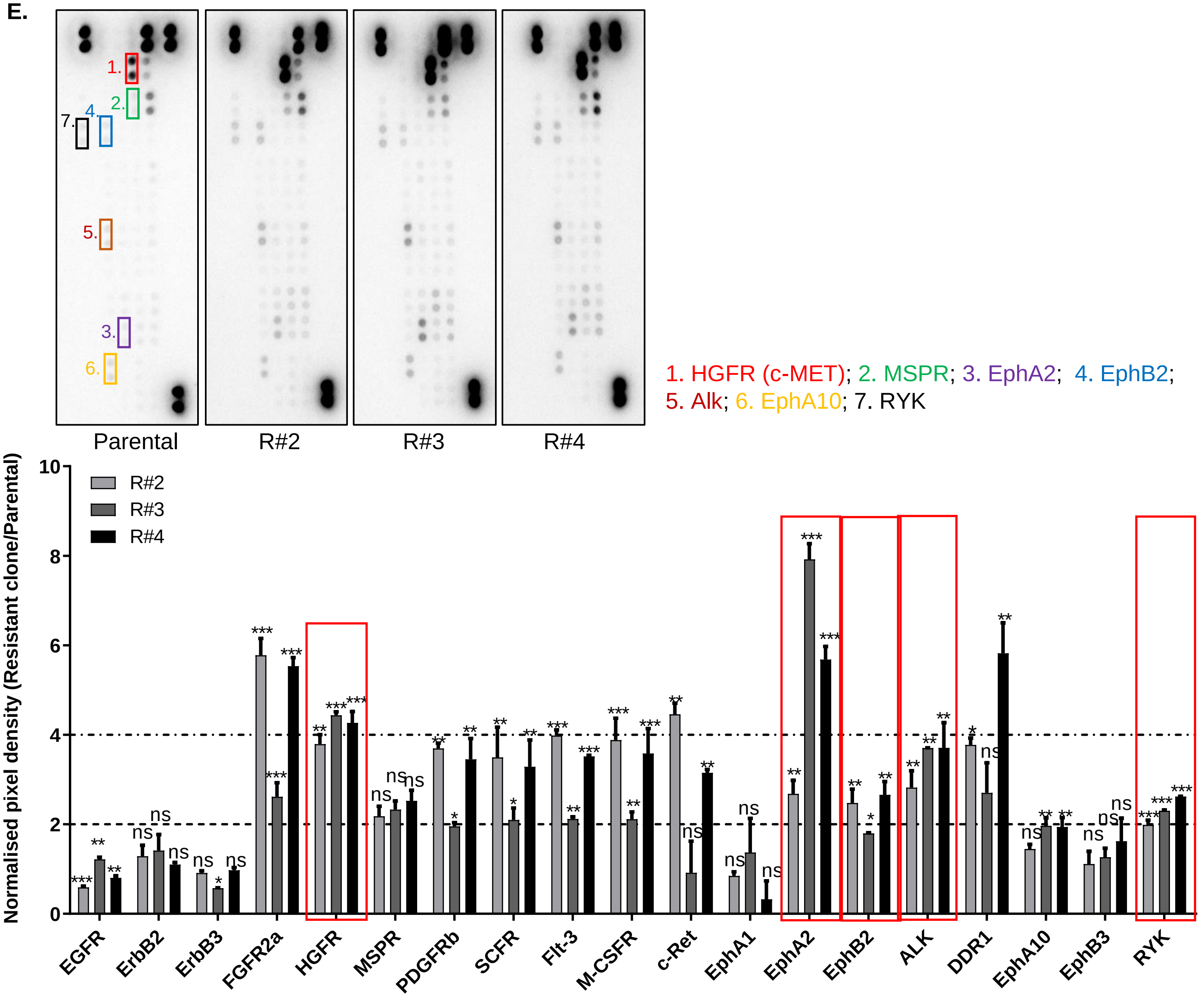
1.
2.
3.
4.
5.
6.
7.
Parental
R#2
R#3
R#4
1. HGFR (c-MET); 2. MSPR; 3. EphA2; 4. EphB2;
5. Alk; 6. EphA10; 7. RYK
Normalised pixel density (Resistant clone/Parental)
R#2
R#3
R#4
EGFR
ErbB2
ErbB3
FGFR2a
HGFR
MSPR
PDGFRb
SCFR
Flt-3
M-CSFR
c-Ret
EphA1
EphA2
EphB2
ALK
DDR1
EphA10
EphB3
RYK

F.

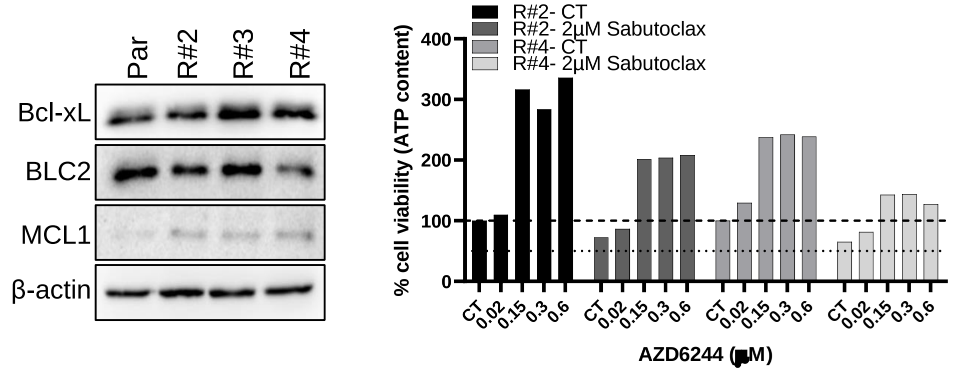
Par
R#2
R#3
R#4
Bcl-xL
BLC2
MCL1
β-actin
R#2- CT
R#2- 2μM Sabutoclax
R#4- CT
R#4- 2μM Sabutoclax
% cell viability (ATP content)
CT
0.02
0.15
0.3
0.6
AZD6244 (μM)

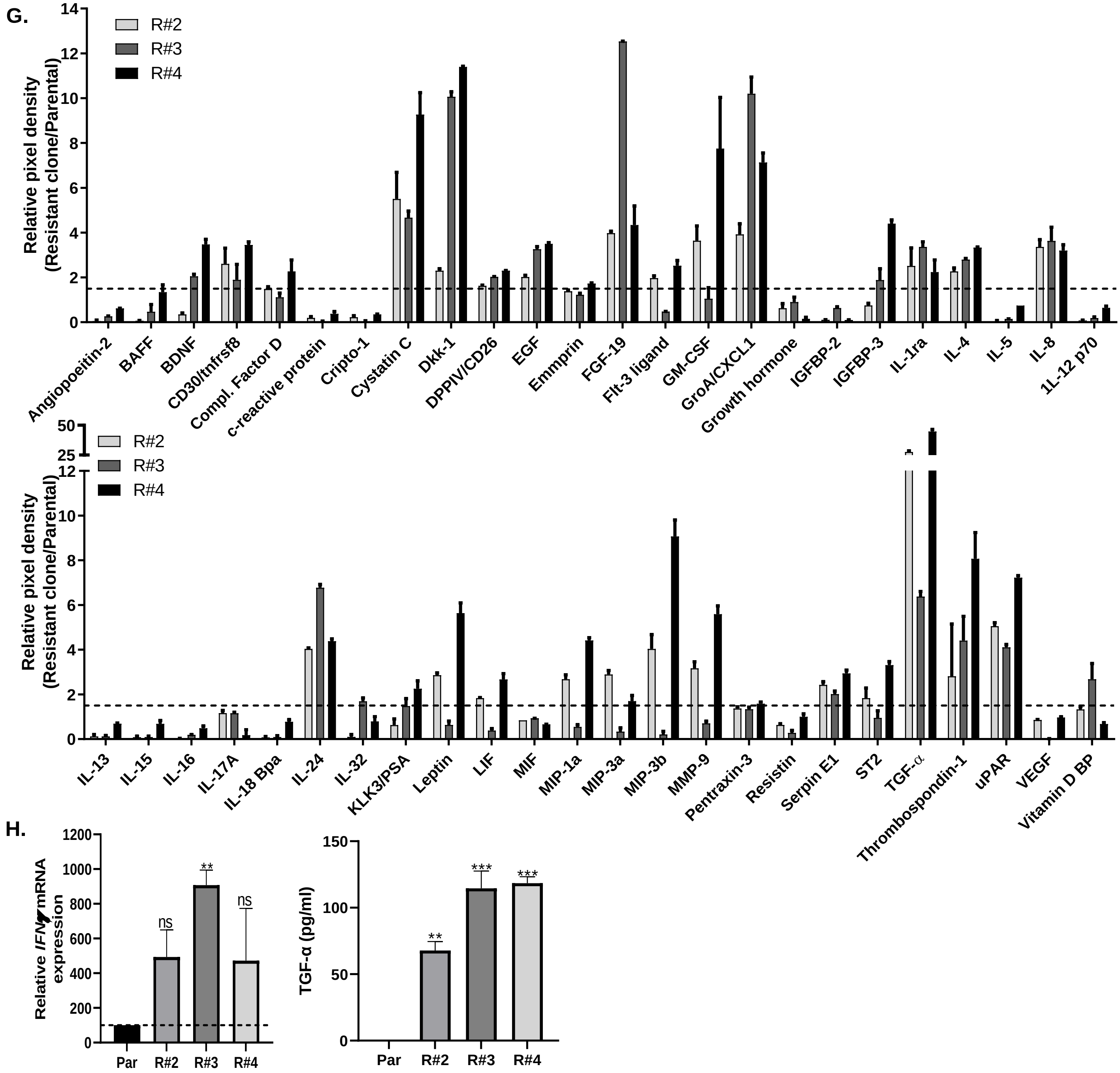


**Supplementary figure 6. Characterization of AZ'1569-R RW7213 cells. A. Left:** Images of morphology of parental and AZ'1569-resistant RW7213 derivatives (#2, #3, #4), obtained using an EVOS microscope (x4 magnification). **Right:** Parental and AZ'1569-R derivatives were treated with MRTX849 for 72h and cell viability determined using CTG assay. $IC_{50}$ values were calculated using Prism software package. **B**. Medexome sequencing of RW7213 parental and AZ'1569-resistant RW7213 clones. Analysis was performed using maftools package in R. Variant classification, type SNV class and top 10 acquired mutated genes for each of the clones are shown. **C**. Results of NGS sequencing of RW7213 Par and AZ'1569-R clones are shown. **D**. **Left panel:** Basal KRAS-GTP and total KRAS expression in RW7213 parental cells, AZ'1569-resistant derivatives and AZ'1569-resistant RW7213 derivatives following removal of AZ'1569 for 8 weeks. **Middle:** pERK1/2, ERK1/2 and KRAS levels in RW7213 parental cells and AZ'1569-resistant derivatives following removal of AZ'1569 for 8 weeks. **Right**: Dose response curve for AZ'1569 in RW7213 parental cells and AZ'1569-R derivatives following removal of AZ'1569 for 8 weeks. **E.** Human phospho-receptor tyrosine kinase array in RW7213 parental and AZ'1569-R clones. The cell extracts were incubated with membranes containing antibodies to 49 different receptor tyrosine kinases. The membranes were washed and incubated with a cocktail of biotinylated detection antibodies to measure the levels of active kinases. Densitometry was performed on the array panels using ImageJ software. **F. Left:** Basal expression levels of Bcl-xL, BCL2 and MCL-1 in RW7213 parental and AZ'1569-resistant clones. **Right:** AZ'1569-resistant RW7213 clone 2 and clone 4 were treated with increasing concentrations of MEK1/2 inhibitor AZD6244 alone or combined with Sabutoclax for 72h and cell viability determined using CTG assay. **G.** Human cytokine array using conditioned medium of RW7213 parental and AZ'1569-resistant clones. Densitometry was performed on the array panels using ImageJ software. **H.** *IFN*$\gamma$ mRNA was quantified using RT-PCR. Raw values were normalised to *ACTB* and *GAPDH* expression and were analysed using the ΔΔCT method. A one-way ANOVA was used to calculate statistical significance. **Right.** TGF-α ELISA using conditioned medium from cells. Data is representative of three independent experimental repeats.
