## Supplementary Materials and Methods for "Bcl-xL is a key mediator of apoptosis following KRAS^G12C^ inhibition in *KRAS*^*G12C*^ mutant colorectal cancer"

#### Cell culture

SNU1411 and LIM2099 were cultured in RPMI-1640, C106 in IMDM, RW7213 in DMEM, SW837 and SW1463 in DMEM/F12 and V481 in MEM2+. Each media was supplemented with 10% FCS and 1mM sodium pyruvate. V481 cells were cultured in MEM2+, supplemented with 1µg/ml hydrocortisone, 10µg/ml Insulin, 2µg/ml Apotransferrin and 0.86ng/ml Sodium Selenite (Merck).

#### DNA expression constructs

Myc-tagged Bcl-xL was purchased from Sino Biological (China). FLAG-MCL-1 (Addgene plasmid#25392) and Flag-BCL2 (Addgene plasmid#18003) were gifts from Roger Davis and Clark Distelhorst, respectively (1).

#### Primers

Primers were purchased from Eurofins Genomics.

<u>RT-PCR</u>

*IFNG* Forward: 5'-TGGCTTTTCAGCTCTGCATC-3', Reverse: 5'-CCGCTACATCTGAATGACCTG -3';

<u>Sanger sequencing</u>

KRAS primers used here have been described in a previous study (2).

*KRAS* exon 2 FW: GGTGGAGTATTTGATAGTGTATTAACC

*KRAS* exon 2 RV: AGAATGGTCCTGCACCAGTAA

*KRAS* exon 3 FW: AAAGGTGCACTGTAATAATCCAGAC

*KRAS* exon 3 RV: ATGCATGGCATTAGCAAAGA

*KRAS* exon 4 FW: TGGACAGGTTTTGAAAGATATTTG

*KRAS* exon 4 RV: ATTAAGAAGCAATGCCCTCTCAAG

**Migration assays**

5µm Transwell polycarbonate membrane inserts from Corning were used. $2.5x10^5$ peripheral blood mononuclear cells (PBMCs) were re-suspended in 200µl of 2% FCS-supplemented DMEM and were added to the top chamber. The bottom chamber was filled with 750µl conditioned medium (2% FCS-supplemented DMEM) obtained from RW7213 parental and resistant cells. Cells were incubated for four hours, following which CellTiter-Glo® was used to measure PBMC migration to the bottom chamber. Serum-free DMEM was used as a negative control. PBMCs were obtained from a healthy donor with written consent and ethical approval granted by the Northern Ireland Blood Transfusion Service, and were provided by Ms. Anne Jordan.

**Immunoprecipitation**

SW837 cells were transfected with 1µg Myc-tagged Bcl-xL for 24 hours, following which cells were treated with AZ'1569, Navitoclax, or combination for a further 24 hours. Cells were lysed with SDS-free RIPA supplemented with protease inhibitors, followed by protein quantification, retrieval of inputs, and incubation with Myc-tag antibody conjugated dynabeads overnight. Dynabeads were then washed and boiled in loading buffer for 5 minutes at 95ºC before beginning the western blotting process.

**Absolute protein quantification for correlation analysis (Log [$EC_{50}$] to Bcl-xL/BAK ratio)**

To calculate BAK and Bcl-xL protein molar concentrations, densitometry values of basal protein expression were obtained for the panel of *KRAS*$^{G12C}$ MT CRC cell lines and HCT116 cells. ImageJ software analysis of western blots for respective proteins, normalised to β-actin (loading control) was used. Fixed molar concentrations were considered for BAK and Bcl-xL in the HCT116 cell line (Bak= 677nM, Bcl-xL = 604nM) as determined previously (3). The equation used was protein (nM) value = ([densitometry value (BAK or Bcl-xL) normalised to β-actin]/respective HCT116 value for BAK or Bcl-xL) x respective concentration in HCT116 (nM). Correlation analyses were performed using GraphPad Prism 9.0.

**Sequencing**

RNA, MedExome and NGS sequencing was performed by the Genomics Core Technology Unit and the Precision Medicine Centre for Excellence, respectively, at Queen's University Belfast.

RNA-sequencing and analysis (GSE198530)

Bulk RNA-sequencing was performed for SW837 and SNU1411 cells treated with 1µM AZ'1569 for 6, 12 and 24 hours. Three biological replicates for each sample were included in the experiment. Total RNA was extracted as described in the main materials and methods section. Each sample underwent QC, and library preparation was performed with 100ng RNA using the KAPA RNA HyperPrep Kit with RiboErase (HMR) (Roche, USA). Libraries were pooled, underwent QC and were sequenced on a NextSeq 500 using a 150 cycle High Output kit (Illumina, USA) to yield >15M paired end 75bp reads per sample.

Generated FASTQ files were analysed using a workflow on Partek® Flow software, v10.0. Briefly, FASTQ files were imported into the software and assigned associated attributes (cell line and time-point), and were aligned with STAR (v2.7.3a) using default settings (assembly=human genome hg38, aligner index=whole genome). Post-alignment QA/QC and quantification of aligned reads to an annotation model (Partek E/M, default settings; min reads=10) was performed, generating Ensembl counts. Differential expression analysis was performed using the GSA (gene specific analysis) tool, with in-built default normalisation applied (normalised to total count, CPM, add 0.0001). 'Time-point' and 'cell line' were chosen as attributes to compare for GSA analysis. The resulting gene lists were downloaded and imported into Ingenuity Pathway Analysis (IPA®) software (Qiagen, UK) to identify significantly enriched pathways for each time-point following AZ'1569 treatment in both cell lines. A cut-off threshold of fold-change >1.3 or <-1.3, and p-value<0.05 was applied to gene lists for pathway analysis. Comparison analysis in IPA® was used to compare significantly enriched ($-log_{10}$ p-value ≥1.3) pathways across both cell lines and all time-points. For volcano plots, $log_2$ratios of time-point vs. control and $-log_{10}$ p-values for genes were plotted on the x- and y-axis respectively, in Prism 9.0.

MedExome sequencing and analysis (PRJNA815942)

DNA was extracted from RW7213 parental and resistant cells as described in the main materials and methods section. Targeted DNA-sequencing was performed with 200ng genomic DNA using the Roche KAPA HyperPlus kit (Roche, USA) for whole genome library preparation and Roche SeqCap EZ MedExome Probes in conjunction with the HyperCap Target Enrichment Kit (Roche) to enrich for targets. This kit enriches coding exons and 20bp of the flanking intronic sequences from around 4600 protein-coding genes, including medically relevant genes (4). Sequencing was performed on the Illumina Novaseq 6000 with a SP 200 cycle reagent kit (Illumina, USA), applying paired end sequencing for 2x 100bp reads.

FASTQ files were aligned to the hg38 genome, sorted, and indexed with bwa mem and samtools (5). Duplicates were marked and all reads assigned to a new read group with Picard (6). Somatic short variants were identified from the samples with a matched normal using GATK (7) (v4.1.9.0) best practices workflow (8, 9). GATK base quality score recalibration was applied, and variants were called with Mutect2. Variants were filtered for contamination and orientation bias artefacts, including standard hard filtering, with FilterMutectCalls as per guidelines. On-target and pass variants were annotated with ANNOVAR (10) for further downstream analysis. VCF files were converted to MAF format using vcf2maf (11) which uses VEP (12). Further analysis and visualisation of MAF files was conducted using maftools (13).

Next Generation Sequencing (NGS) and analysis

DNA was extracted from RW7213 parental and AZ'1569-resistant cells as described in the main methods section. 200ng genomic DNA was used as the input for NGS libraries using the KAPA HyperPlus Kit (Roche Sequencing Solutions, Pleasanton, CA, US) according to the manufacturer's instructions and in-house SOPs. Prepared DNA libraries were hybridised with a custom-designed hybrid-capture next generation sequencing panel (Roche Sequencing

Solutions). Following hybridisation, samples were sequenced on a NextSeq 500 (Illumina, San Diego, CA, USA) using the NextSeq 550 System Mid-Output Kit v2 (150 cycles) with 76bp paired-end reads. Binary base call (BCL) files were processed using custom-designed in-house bioinformatics pipelines which converted BCL files to FASTQ files, in accordance with Genome Analysis Toolkit (GATK) best practices (7), including alignment to the hg38 build of the human reference genome using the Burrows-Wheeler Alignment (BWA) tool. Following deduplication and quality scoring, variant calling was performed using Vardict (14) for SNP/INDEL variant calling, Manta for structural variant calling (15), CNVPanelizer (16) for copy number estimation and MSIsensor-Pro (17) for microsatellite instability assessment. Variant filtering and annotation was performed using SnpEff (18) and customised scripts developed specifically for this assay.

<u>Data availability</u>

Raw data for RNA-sequencing and MedExome sequencing experiments has been deposited at the relevant NCBI platforms, under the accession numbers GSE198530 and PRJNA815942, respectively.

#### ***In vivo* study**

<u>Growth curve studies</u>

Tumour growth was assessed following injection of 2.5 $x10^6$ and $5x10^6$ SNU1411, 7.5 $x10^6$ and $10x10^6$ SW837 and $7.5x10^6$ and $10x10^6$ SW1463 cells in the left and right flank of mice respectively. Mice weight and xenograft growth (using the formula: *(shortest tumour diameter)*$^2$ × *longest tumour diameter* × *0.5*) was monitored 3 times per week. Mice were sacrificed when combined tumour volume reached $1200mm^3$ or 1 month following cell inoculation. In contrast to the SW837 model, SNU1411 and SW1463 xenografts both showed exponential growth (Supplementary Fig. S5A) and hence were taken forward in the efficacy study.

<u>Tolerability study</u>

The maximum tolerated dose of $KRAS^{G12C}$ inhibitor AZ’8037 (oral gavage; 50 or 100mg/kg) in combination with Navitoclax (oral gavage; 50, 75 or 100mg/kg), was assessed in NOD/SCID mice in a dose-escalation study. Navitoclax was formulated in 10% ethanol, 30% polyethylene glycol 400, and 60% Phosal 50 PG and AZ’8037 in 1% (w/v) Pluronic F127. A single dose of the lowest dose of AZ’8037 50mg/kg PO (oral gavage), was combined with Navitoclax (50mg/kg). Doses were increased if no adverse events were noticed after 2 treatments with an interval of three days (to explore late toxicity) on the lower dose. Outward signs of distress and mouse weight were monitored daily was monitored. MTD was defined as the maximal dose of both drugs in combination which does not results in weight loss > 15% or death (Supplementary Fig. S5B).

### REFERENCES


1. Wang NS, Unkila MT, Reineks EZ,Distelhorst CW. Transient expression of wild-type or mitochondrially targeted Bcl-2 induces apoptosis, whereas transient expression of endoplasmic reticulum-targeted Bcl-2 is protective against Bax-induced cell death. *J Biol Chem* 2001; 276: 44117-28.
2. Oddo D, Sennott EM, Barault L, Valtorta E, Arena S, Cassingena A , et al. Molecular Landscape of Acquired Resistance to Targeted Therapy Combinations in BRAF-Mutant Colorectal Cancer. *Cancer Res* 2016; 76: 4504-15.
3. Lindner AU, Concannon CG, Boukes GJ, Cannon MD, Llambi F, Ryan D , et al. Systems analysis of BCL2 protein family interactions establishes a model to predict responses to chemotherapy. *Cancer Res* 2013; 73: 519-28.
4. Aref-Eshghi E, Kerkhof J, Carere DA, Volodarsky M, Bhai P, Colaiacovo S , et al. Clinical and technical assessment of MedExome vs. NGS panels in patients with suspected genetic disorders in Southwestern Ontario. *J Hum Genet* 2021; 66: 451-64.
5. Li H, Handsaker B, Wysoker A, Fennell T, Ruan J, Homer N , et al. The Sequence Alignment/Map format and SAMtools. *Bioinformatics* 2009; 25: 2078-9.
6. Picard Toolkit. [Internet]. Broad Institute, GitHub Repository.; Available from: https://broadinstitute.github.io/picard/.
7. McKenna A, Hanna M, Banks E, Sivachenko A, Cibulskis K, Kernytsky A , et al. The Genome Analysis Toolkit: a MapReduce framework for analyzing next-generation DNA sequencing data. *Genome research* 2010; 20: 1297-303.
8. DePristo MA, Banks E, Poplin R, Garimella KV, Maguire JR, Hartl C , et al. A framework for variation discovery and genotyping using next-generation DNA sequencing data. *Nat Genet* 2011; 43: 491-8.
9. Van der Auwera GA, Carneiro MO, Hartl C, Poplin R, Del Angel G, Levy-Moonshine A , et al. From FastQ data to high confidence variant calls: the Genome Analysis Toolkit best practices pipeline. *Curr Protoc Bioinformatics* 2013; 43: 11 0 1- 0 33.
10. Wang K, Li M,Hakonarson H. ANNOVAR: functional annotation of genetic variants from high-throughput sequencing data. *Nucleic acids research* 2010; 38: e164.
11. Kandoth C. mskcc/vcf2maf: vcf2maf. .
12. McLaren W, Gil L, Hunt SE, Riat HS, Ritchie GR, Thormann A , et al. The Ensembl Variant Effect Predictor. *Genome Biol* 2016; 17: 122.
13. Mayakonda A, Lin DC, Assenov Y, Plass C,Koeffler HP. Maftools: efficient and comprehensive analysis of somatic variants in cancer. *Genome research* 2018; 28: 1747-56.

14. Lai Z, Markovets A, Ahdesmaki M, Chapman B, Hofmann O, McEwen R , et al. VarDict: a novel and versatile variant caller for next-generation sequencing in cancer research. *Nucleic acids research* 2016; 44: e108.
15. Chen X, Schulz-Trieglaff O, Shaw R, Barnes B, Schlesinger F, Kallberg M , et al. Manta: rapid detection of structural variants and indels for germline and cancer sequencing applications. *Bioinformatics* 2016; 32: 1220-2.
16. Cristiano Oliveira, Thomas Wolf: CNVPanelizer: Reliable CNV detection in targeted sequencing applications. R package version 1.22.0. (2020). https://bioconductor.org/packages/release/bioc/html/CNVPanelizer.html.
17. MSIsensor-pro: Fast, Accurate, and Matched-normal-sample-free Detection of Microsatellite Instability. https://doi.org/10.1016/j.gpb.2020.02.001.
18. A program for annotating and predicting the effects of single nucleotide polymorphisms, SnpEff. doi: 10.4161/fly.19695.
