## Supplementary Tables 1 and 2 for "Bcl-xL is a key mediator of apoptosis following KRAS^G12C^ inhibition in *KRAS*^*G12C*^ mutant colorectal cancer"

Supplementary Tables 1&2

### Supplementary Table 1: Antibodies

| Antibody | Supplier | Product code | Animal | Dilution |
|---|---|---|---|---|
| Anti-mouse HRP | Cell Signalling Technology | 7076 | Horse | 1: 2000 |
| Anti-Rabbit HRP | Cell Signalling Technology | 7074 | Goat | 1: 2000 |
| Bad | Cell Signalling Technology | 9239 | Rabbit | 1: 1000 |
| Bax | Cell Signalling Technology | 5023 | Rabbit | 1: 1000 |
| Bcl-2 | Cell Signalling Technology | 2872 | Rabbit | 1: 1000 |
| Bid | Cell Signalling Technology | 2002 | Rabbit | 1: 1000 |
| Bak | Cell Signalling Technology | 12105 | Rabbit | 1: 1000 |
| Bim | Cell Signalling Technology | 2933 | Rabbit | 1: 1000 |
| Bcl-xL (54H6) | Cell Signalling Technology | 2764 | Rabbit | 1: 1000 |
| Cleaved Caspase-3 | Cell Signalling Technology | 9661 | Rabbit | 1: 500 |
| Caspase-8 (12F5) | Enzo | ALX-804-242-C100 | Mouse | 1: 2000 |
| Caspase-9 | Cell Signalling Technology | 9502 | Rabbit | 1: 1000 |
| Mcl-1 | BD Transduction | 559027 | Mouse | 1: 1000 |
| PARP | Cell Signalling Technology | 9542 | Rabbit | 1: 2000 |
| KRAS | LS-Biosciences | LS-C175665 | Mouse | 1: 2000 |
| pEGFR (Tyr1068) | Sigma-Aldrich | 324867 | Rabbit | 1: 1000 |
| EGFR | BD Transduction | 89414 | Mouse | 1: 1000 |
| pERK1/2 (Thr202/Tyr204) | Cell Signalling Technology | 9101 | Rabbit | 1: 1000 |
| ERK1/2 | Cell Signalling Technology | 9102 | Rabbit | 1: 1000 |
| pMEK1/2 (Ser217/221) | Cell Signalling Technology | 9154 | Rabbit | 1: 1000 |
| MEK1/2 | Cell Signalling Technology | 9122 | Rabbit | 1: 1000 |
| pS6 (Ser235/236) | Cell Signalling Technology | 4857 | Rabbit | 1: 1000 |
| S6 | Cell Signalling Technology | 2317 | Mouse | 1: 1000 |
| pAKT (Ser 473) | Cell Signalling Technology | 4060 | Rabbit | 1: 1000 |
| AKT | Cell Signalling Technology | 9272 | Rabbit | 1: 1000 |
| pMET (Tyr1234/1235) | Cell Signalling Technology | 3077 | Rabbit | 1: 1000 |
| MET | Cell Signalling Technology | 8198 | Rabbit | 1: 1000 |
| pEphA2 (Ser897) | Cell Signalling Technology | 6347 | Rabbit | 1: 1000 |
| pEphA2 (Tyr588) | Cell Signalling Technology | 12677 | Rabbit | 1: 1000 |
| pEphA2(Tyr772) | Cell Signalling Technology | 8244 | Rabbit | 1: 1000 |
| EphA2 | Cell Signalling Technology | 12927 | Mouse | 1: 1000 |
| FLAG (M2) HRP | Sigma | A8592 | Mouse | 1: 10,000 |
| Myc-Tag (71D10) | Cell Signalling Technology | 2278 | Rabbit | 1: 1000 |
| β-actin (AC-74) | Sigma | A2228 | Mouse | 1: 10,000 |
| All antibodies were made up in 5% milk (Marvel,UK)/PBS-Tween | | | | |

### Supplementary Table 2: Compound concentrations for screen.

| Product Name | Pathway | Target | Concentration (µM) | Stock concentration (µM) |
|---|---|---|---|---|
| ABT-737 | Apoptosis/cell death | Bcl-2,Autophagy | 0.5, 1, 2 | 10mM |
| Saracatinib (AZD0530) | Angiogenesis/ Src inhibitor | Src | 0.25, 0.5, 1 | 10mM |
| Vorinostat (SAHA, MK0683) | Epigenetics | Autophagy,HDAC | 0.5, 1, 2 | 10mM |
| Entinostat (MS-275) | Epigenetics | HDAC | 0.5, 1, 2.5 | 10mM |
| Olaparib (AZD2281, Ku-0059436) | DNA damage | PARP | 1, 2, 3 | 10mM |
| Nutlin-3 | MDM2 antagonist/ p53/apoptosis | E3 Ligase ,Mdm2 | 1, 5, 10 | 10mM |
| Vismodegib (GDC-0449) | Wnt signalling pathway | Hedgehog/Smoothened | 1, 5, 10 | 10mM |
| Alisertib (MLN8237) | Cell cycle | Aurora Kinase | 0.03, 0.125, 0.5 | 10mM |
| Barasertib (AZD1152-HQPA) | Cell cycle | Aurora Kinase | 0.02, 0.1, 0.5 | 10mM |
| Roscovitine (Seliciclib,CYC202) | Cell cycle | CDK | 2.5, 5, 10 | 10mM |
| Ganetespib (STA-9090) | cytoskeletal signalling/ HSP90 inhibitor | HSP (e.g. HSP90) | 0.25, 0.5, 1 | 10mM |
| BIBR 1532 | DNA damage | Telomerase | 5, 10, 20 | 10mM |
| Epothilone A | cytoskeletal signalling | Microtubule Associated | 0.002, 0.02, 0.2 | 10mM |
| AZD7762 | cell cycle | Chk | 0.125, 0.25, 0.5 | 10mM |
| Ixazomib (MLN2238) | proteosome inhibitor | Proteasome | 0.1, 0.25, 0.5 | 10mM |
| Degrasyn (WP1130) | proteosome inhibitor | Bcr-Abl,DUB | 1, 1.5, 2 | 10mM |
| Rosiglitazone | Metabolism | PPAR | 0.1, 1, 10 | 10mM |
| AT406 (SM-406) | Cell death/IAP | E3 Ligase ,IAP | 1, 2.5, 5 | 10mM |
| I-BET151 (GSK1210151A) | Epigenetics | Epigenetic Reader Do | 0.5, 1, 2 | 10mM |
| Sirtinol | Epigenetics | Sirtuin | 5, 10, 20 | 10mM |
| Carfilzomib (PR-171) | proteosome inhibitor | Proteasome | 0.01, 0.02, 0.04 | 10mM |
| IMD 0354 | NF-kB | IκB/IKK | 0.5, 1, 2.5 | 10mM |
| Salubrinal | ER stress/ UPR | PERK | 5, 10, 20 | 10mM |
| JNK-IN-8 | MAPK pathway | JNK | 1, 3, 5 | 10mM |
| Birinapant | IAP inhibitor/cell death | IAP | 0.1, 1, 10 | 10mM |
| RG-7112 | Apoptosis/Mdm2 inhibitor | Mdm2 | 0.1, 1, 2.5 | 10mM |
| AZD1208 | JAK/STAT | Pim | 1, 3, 5 | 10mM |
| UNC1999 | Epigenetics | Histone Methyltransferase | 1, 2, 3 | 10mM |
| Tasisulam | caspase activator/cell death/ | Caspase | 1, 3, 10 | 10mM |
| TH287 | DNA damage | MTH1 | 0.5, 1, 3 | 10mM |
| AZD6738 | DNA damage | ATM/ATR | 0.5, 1, 3 | 10mM |
| Venetoclax (ABT-199, GDC-0199) | Apoptosis | Bcl-2 | 0.5, 1, 3 | 10mM |
| Sabutoclax | Apoptosis | Bcl-2, Bcl-xl, Mcl-1, Bfl-1 | 0.25, 1, 2.5 | 10mM |
| CB-5083 | Transmembrane transporters | ATPase | 0.25, 0.5, 1 | 10mM |
| AZD1390 | DNA damage | ATM/ATR | 0.01, 0.1, 0.3 | 10mM |
| Palbociclib (PD-0332991) HCl | Cell cycle inhibitor | CDK | 0.5, 1, 2.5 | 10mM |
| CX-5461 | DNA damage | DNA/RNA Synthesis | 0.1, 0.3, 1 | 2mM |
| NU7026 | DNA damage | DNA-PK | 2, 5, 10 | 2mM |
| Palifosfamide | DNA damage | | 1, 2.5, 5 | 10mM |
| EPZ5676 | Epigenetics | DOT1L histone methyltransferase | 0.5, 1, 3 | 10mM |
| AZD5991 | Cell death- Mcl-1 inhibitor | Bcl2 | 0.25, 1, 2.5 | 10mM |
| AZD4573 | Cell cycle | CDK9 | 0.1, 0.3, 1 | 10mM |
| Durvalumab | Immunology & Inflammation | PDL1 | 0.2, 2, 20 | 10mM |
| ONC206 | ER stress/ UPR | DRD2 | 0.25, 0.5, 1 | 10mM |
| iz-TRAIL | Cell death | DR4/5 | 0.5, 1, 2ng/ml | 1.93mg/ml |
